# Direct observation of Notch signalling induced transcription hubs mediating gene-expression responses

**DOI:** 10.1101/2025.07.04.663121

**Authors:** Carmen Santa-Cruz Mateos, Charalambos Roussos, F. Javier de Haro Arbona, Julia Falo-Sanjuan, Sarah Bray

## Abstract

Developmental decisions rely on cells making accurate transcriptional responses to signals they receive, as with Notch pathway activity. Local condensates or transcription factor hubs are a proposed mechanism for facilitating gene activation by nuclear complexes. To investigate their importance in endogenous Notch signalling, we deployed multi-colour live-imaging to measure Notch transcription-complex enrichment at a target gene locus in combination with the transcription dynamics. The co-activator Mastermind (Mam) was present in signalling-dependent nuclear foci, during Notch active developmental stages. Tracking these highly-dynamic Mam hubs together with transcription in the same nucleus, revealed that their condensation precedes and correlates with the profile of transcription and becomes stabilized if transcription is inhibited. Manipulations to signalling levels had concordant effects on hub intensities and transcription profiles, altering their probability and amplitude. Together the results argue that signalling induces the formation of transcription hubs whose properties are instrumental in the quantitative gene expression response to Notch activation.

## INTRODUCTION

To make and organise different tissues, cells decipher information from developmental signalling pathways. Transmitting this information accurately, so that cell-surface signals are translated into correct transcriptional responses, is of critical importance. For example, dosage and dynamics of Notch activity are fundamentally important for developmental decisions, for sustaining stem cells and for tissue homeostasis ^1–5^. Misregulation underlies many diseases including cancers ^6–8^. Signalling is normally initiated when transmembrane ligands on adjacent cells interact with the Notch receptor at the membrane. This brings about proteolytic cleavages to release the intracellular domain, NICD, which forms a tripartite complex with the DNA binding protein CSL and coactivator Mastermind ^9–11^. The tripartite complex is recruited directly to regulated gene loci, where it promotes their transcription^12–14^. As there is no amplification step, the numbers of active transcription complexes directly relate to the number of receptors activated and it remains to be established how they efficiently and quantitatively elicit target gene transcription.

One model is that Notch transcription complexes form hubs, local zones of high-density clustering that facilitate the recruitment of transcription machinery and RNA Polymerase II (Pol II) to direct transcription ^15,16^. Hubs, or condensates, have been detected in a range of contexts where their formation relies on protein-DNA and multivalent protein-protein interactions, often involving low-complexity regions ^17–20^. Broadly, they can be described as non-stoichiometric assemblies, which drive biological functions by transiently increasing local concentrations ^21,22^. Although initially detected under conditions of protein over-expression transcription factor hubs are increasingly observed in physiological conditions making it plausible that they perform an important step in transcriptional regulation ^23–26^. Indeed, recent studies suggest that hub densities correlate with the transcriptional output from gene loci ^27^.

There is mounting evidence that condensation of nuclear hormone receptors and coregulators play an important role in transcriptional regulation ^28,29^. Assembly of nuclear condensates containing yes-associated protein (YAP) and transcription coactivators has also been seen in cells, where it is modulated by Hippo signalling or mechanical changes^30^. Similarly, using live imaging approaches, we previously observed a hub-like enrichment of Notch transcription complexes around a target locus, in a tissue where ectopic Notch activity was induced ^15,16^. As with the YAP condensates, the zone of enrichment concentrated additional factors that included Mediator and Pol II in those nuclei where transcription was occurring ^15,30^. It is thus plausible that such hubs play an integral part in signalling responses.

To explore the role of transcription hubs in Notch signalling we focus here on a biological context where the Notch pathway is endogenously active, the *Drosophila* follicular epithelium. Through in vivo live imaging of endogenous members of Notch transcription complexes together with a method for labelling the Notch-regulated *E(spl)*-Complex [*E(spl)-C*] we could detect localised condensates or transcription hubs in the stages when Notch signalling is occurring. To investigate the role of these hubs in mediating the response to Notch, we tracked the co-activator Mam, a component of the Notch active transcription complexes, at the same time as measuring responding gene expression profiles using the MS2/MCP system with two Notch targets. This revealed that the Mam transcription hubs are dynamic and reach their maximal intensity just prior to transcription initiation. Hub intensity is altered by changes in Notch activity, indicating that they are signalling induced, and the consequential effects on transcription argue that gene-specific Mam transcription hubs have a key role in coordinating the probability and levels of gene activity in response to Notch activation.

## RESULTS

### Mastermind hubs are present during Notch active time-window

The active Notch transcription complex, containing Mastermind (Mam), forms a local hub of enrichment at a target gene complex under conditions of prolonged ectopic activity ^15^. To investigate whether similar hubs occur when the pathway is endogenously active, we chose to focus on the follicular epithelium (FE) in the *Drosophila* ovary which undergoes a period of Notch signalling between stages 5 to 8 of egg chamber development ^31^. Notch activity in this monolayer follicular epithelium relies on ligands produced by the underlying germ-line cells ^32,33^, as illustrated by the expression of Delta:mScarlet, which is present in the membranes at the interface between the germline and the follicle cells (Figure 1A,B). The broad distribution of Delta is consistent with the adjacent FE cells having an equivalent potential for Notch activity.

**Figure 1:**
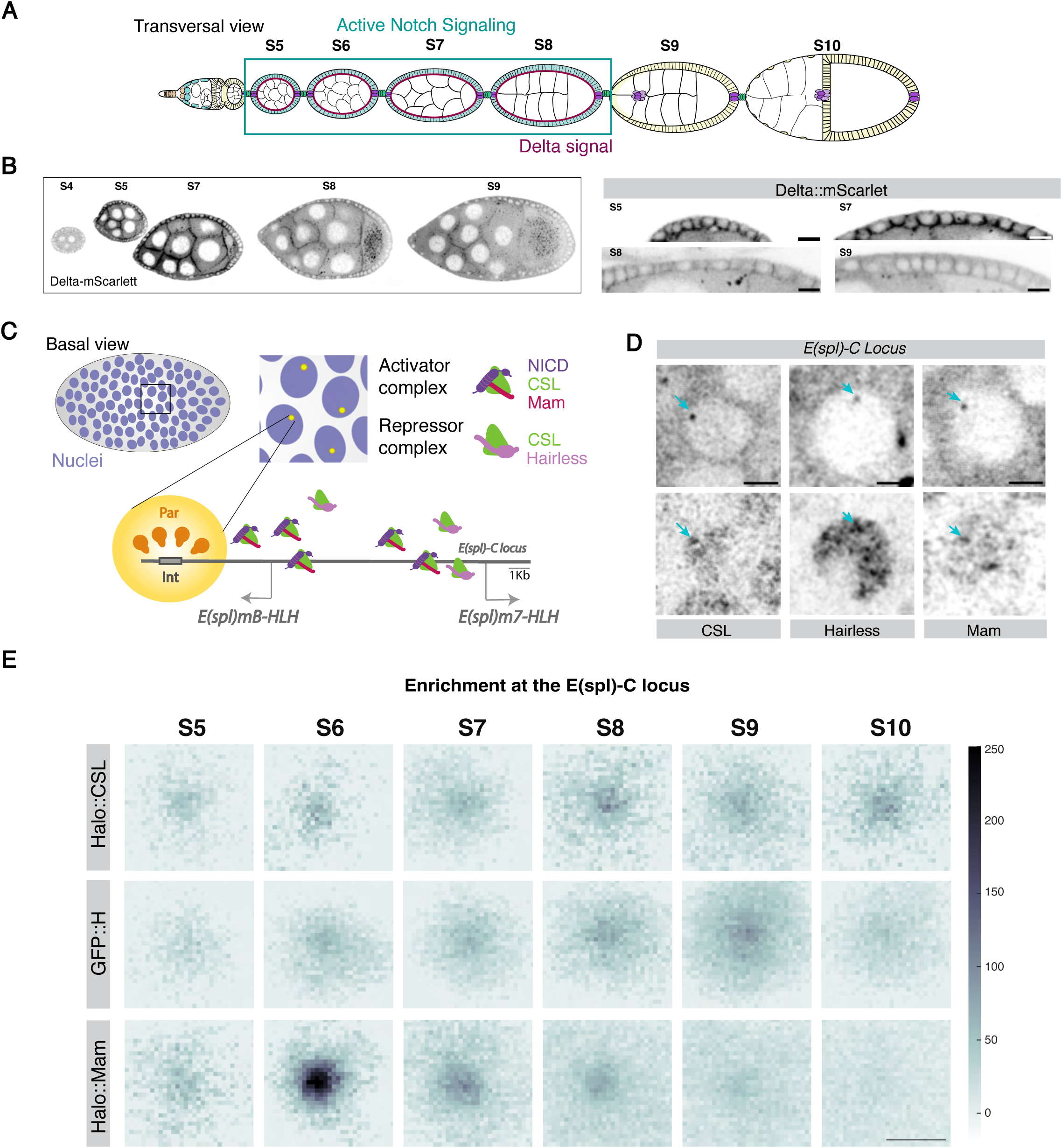
Mastermind hub forms at *E(spl)-C* during stages with endogenous Notch activity. (A) Cartoon of a *Drosophila* ovariole. The follicular epithelium (FE, pale blue, yellow) is a simple epithelium that surrounds the germline (uncoloured). Notch is active between stages 5 and 9 of development (blue rectangle) due to expression of Delta (magenta) on the surface of germline cells during those stages. (B) Confocal images from Delta-mScarlet showing expression between stages 4 and 9 (right) with higher magnification images of the membranes at FE-germline interface at the stages indicated. Scale bars represent 10 μm. (C) Diagram summarising the method using Par/Int to label *E(spl)-C*. In follicle cell (FC) nuclei (purple), *E(spl)-C* is detected through binding of Par protein (yellow) to the *Int* sequence (grey) inserted within the endogenous locus. Recruitment at the locus of activator complexes (NICD [purple], CSL [green] and Mastermind [Magenta]) or co-repressor complexes (CSL [green], Hairless [light pink]) can be measured. (D) Live imaging of individual FC nuclei showing the location of the *E(spl)-C* locus (upper panels) and the localised enrichment of Halo::CSL, GFP::Hairless or Halo::Mam (lower panels). Scale bars represent 2μm. (E) Average enrichment of the indicated proteins at *E(spl)-C* normalized to a random nuclear region (see methods). CSL and Hairless are recruited in all stages whereas Mam is enriched from stage 5 to 8 coincident with Notch-active time window. Number of analyzed nuclei for Halo::CSL (221, 137, 377, 137, 259, 158), Halo::Mam (92, 141, 173, 171, 287, 219) and GFP::Hairless (94,186, 241,158, 248, 207) were from at least 3 e.c. per condition. Scale bar represents 1 μm.

To investigate the formation of transcription hubs in this context, we analysed the distribution of endogenously tagged factors by live-imaging ^15^. The core transcription factor CSL and its partners the coactivator Mam and the corepressor Hairless are all present in follicular cell nuclei throughout the period analysed (Supplementary Figure S1A). All exhibited a degree of heterogeneity within each nucleus, consistent with them forming condensates or “hubs”. To investigate if these hubs are associated with Notch regulated loci, we took advantage of the Par/Int system to identify the *Enhancer of split complex [E(spl)-C]* locus in live tissues ^15,16,34^. This relies on recruitment of fluorescently tagged Par binding proteins to their cognate *int* sequences which have been engineered into the *E(spl)-C* locus and result in a locus-specific fluorescent spot within each nucleus (Figure 1C,D). The levels of factors colocalising with this tagged locus were then compared to those elsewhere in the nucleus. Results from multiple nuclei were pooled and the average intensity centered on the locus region was visualised by heatmaps (Figure 1E, Supplementary Figure S1B).

All three members of Notch transcription complexes CSL, Mam and Hairless were significantly enriched at the *E(spl)-C* locus during the stages analysed (Figure 1D,E) whereas an unrelated transcription factor, Sox 14, was not (Supplementary Figure S1C). The DNA-binding CSL was enriched from stages 5 to 10, and at broadly equivalent levels from stage 6 onwards (Figure 1E). In contrast, the coactivator Mam was enriched in a more restricted time window, from stage 5-8 with the highest levels detected during stage 6 (Figure 1E). The period of Mam recruitment corresponds to the time when the Notch pathway is active ^31,35^ and the locus-specific enrichment of Mam was absent in tissues exposed to a gamma-secretase inhibitor for 4 hours to block the activating cleavage (Supplementary Figure 1D). These data demonstrate that Mam hub formation requires Notch activity. The average diameter of the Mam puncta is circa 500 nm which is close to the diffraction limit of the microscope and the majority may be smaller based on estimates from single molecule tracking ^36^. Overall, they have similar properties to the hubs formed by the Bicoid and Dorsal transcription factors in the embryo ^27,37,38^ suggesting they correspond to similar transcription factor condensates.

In the absence of Notch activity, CSL is associated with the co-repressor Hairless and, in agreement, Hairless was associated with the target locus during a more prolonged period and became more highly enriched in stages 9-10 (Figure 1E). The fact that CSL and Hairless remain enriched after Notch is no longer active (stages 9 and 10) is consistent with a repressive role in those later stages and with evidence indicating that regulated loci retain a memory of Notch activity for several hours ^15^. Furthermore, the presence of all 3 proteins during the period of Notch pathway activity fits with the proposal that dynamic exchange between activator and corepressor complexes tunes the response to signalling ^9,15^. Here we primarily focus on the striking recruitment of the co-activator Mam as a measure of a signalling-induced transcription hub at an endogenous locus.

### Dynamics of Notch target-gene transcription in follicular cells

The live imaging indicates that Mam transcription complexes are enriched at *E(spl)-C* during the period of endogenous Notch activity. To investigate the relationship between the Mam hubs and target gene transcription, we began by characterising the transcriptional response in the FE using single-molecule fluorescence in situ hybridization (smFISH)^39,40^. Using fluorescently labelled probes for *E(spl)m7* and *E(spl)m*β, both of which are expressed in the FE in response to Notch signalling ^35,41,42^, we focused our analysis on the bright nuclear foci that correspond to the Active Transcription Sites (ATS) as a measure of the actively transcribing cells at each time point (Robinson-Thiewes et al., 2020; Figure 2A). Only one ATS was detected per nucleus for each gene and contrary to expectations, only a low proportion of nuclei contained an ATS for one or both genes at any given time point (Figure 2B, Supplementary Figure S2A). More nuclei contained ATS for *E(spl)-m*β, (15-20% at stage 6) than for *E(spl)m7* (4-5% at stage 6) (Figure 2B, Supplementary Figure S2A). However, the majority (>90%) of *E(spl)m7* ATS co-localised with an *E(spl)-m*β ATS suggesting they respond to the same signalling input, and that E(spl)*m*β has a higher probability of transcription than E(spl)*m7*.

**Figure 2:**
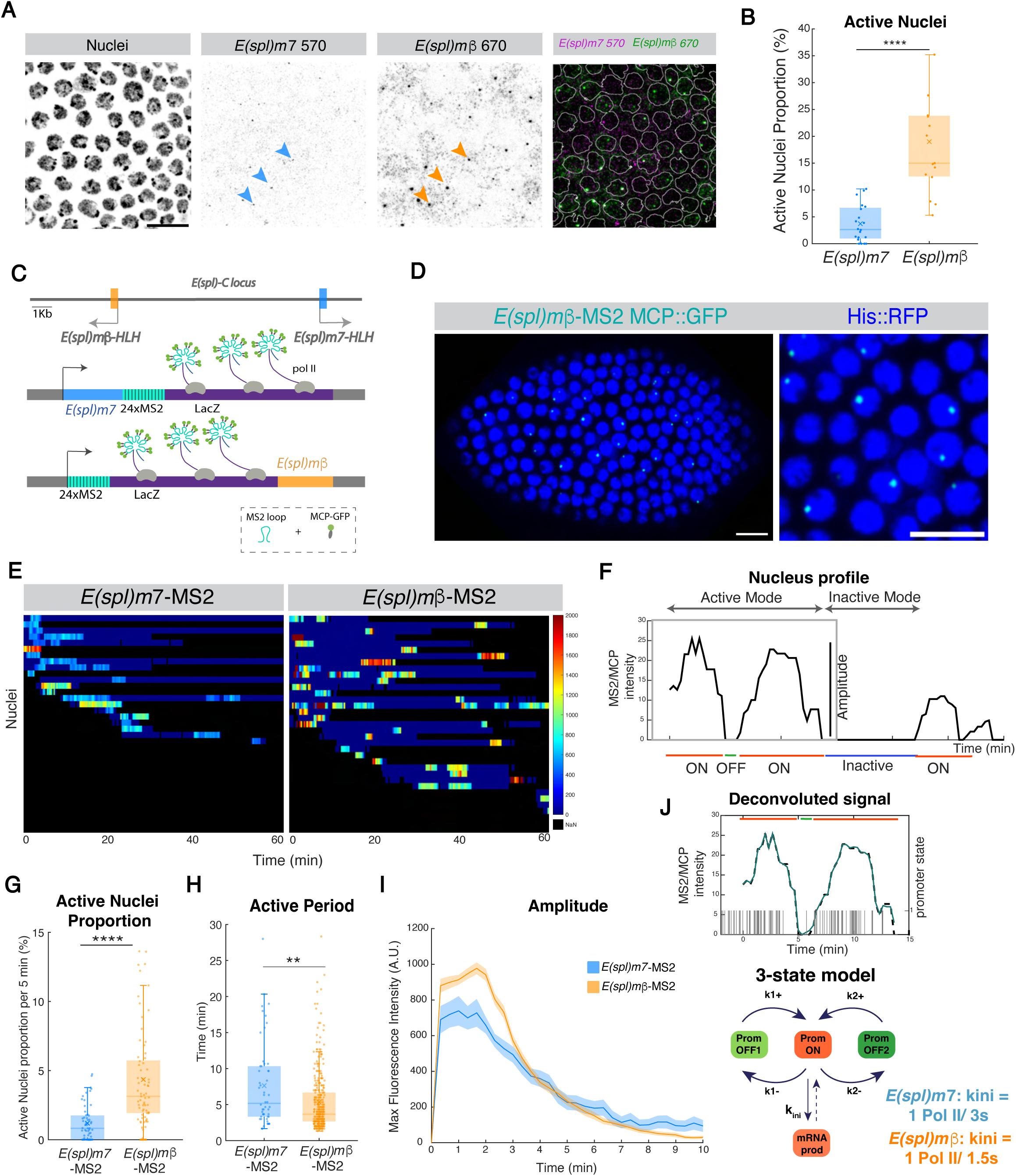
Transcription of Notch responsive genes is probabilistic and pulsatile. (A) Confocal images of smFISH with *E(spl)-m7*-570 and *E(spl)-m*β-670 probes in stage 6 egg chambers (e.c.). Active transcription sites (ATS) for *E(spl)-m7* (blue arrowheads) and *E(spl)-m*β (orange arrowheads) colocalize. Scale bar represents 10 μm. (B) Proportion of ATS for *E(spl)m7* and *E(spl)m*β at Stage 6 detected by smFISH. n= 20 e.c. (*m7*) and 18 e.c. (*m*β). (C) Diagram illustrating the MS2/MCP system. 24x MS2 loops (cyan) were introduced into the endogenous *E(spl)m7 (blue,* coding region) and E(spl)*m*β (orange, coding region) genes along with lacZ (purple) to extend the length of the RNA produced aiding detection. When transcribed (PolII, grey), MCP-GFP (green) binds to the MS2 loops in the nascent RNA. (D) Image from *in vivo* movie of *E(spl)m*β*-MS2,* transcription is detected as puncta of MCP-GFP (cyan) within nuclei (blue, H2Av-RFP). Scale bars represent 10 μm. (E) Kymograph heatmap of maximum fluorescence intensity in active nuclei with time from representative movies of *E(spl)m7*-MS2 and *E(spl)m*β*-MS2* at stage 6. (Supplementary movies S1, S2) (F) Example of transcription trace from a single nucleus with active and inactive periods and promoter states indicated. (G) Average active nuclei proportion per 5-minute period for *E(spl)m7-MS2* (blue) and *E(spl)m*β*-*MS2 (orange) at stage 6. p = 3.16e-9. (H) Active period durations for *E(spl)m7-MS2* (blue) and *E(spl)m*β*-*MS2 (orange) at stage 6. p = 0.0018. G-H Boxplots indicate median, with 25–75 quartiles; error bars are SD. (I) Average intensities of transcription traces from *E(spl)m7*-MS2 and *E(spl)m*β-MS2 at stage 6. Shade surrounding average line represents standard error of the mean (SEM); n = 66 nuclei from 6 e.c. for *E(spl)m7*-MS2, n=443 nuclei from 6 e.c. for *E(spl)m*β -MS2. (J) Example MS2 trace of a particular nuclei through time. Intensity of MCP is showed together with the promoter state calculated using BURSTDECOV model to deconvolute the signal (see Material and Methods). For both genes, 3-state model, with one ON state and two OFFs, is the minimal number of states that explained data. Major difference between *E(spl)m7*-MS2 and *E(spl)m*β*-*MS2 is kini.

Given the requirement for Notch activity throughout the epithelium and the duration of the developmental stage ^43^, all nuclei are likely to undergo one or more periods of transcriptional activity. The relatively low proportion of nuclei with ATS at any one time-point suggests that each nucleus transcribes for short “active” periods, interspersed with longer periods of inactivity. To investigate this possibility and further analyse the dynamics of transcription at stage 6 we employed the MS2/MCP system, to image transcription from *E(spl)m7* and *E(spl)m*β in real time ^44^. MS2 loops were introduced into the endogenous genes by CRISPR-Cas9 engineering (see methods) and the resulting strains combined with MCP::GFP and Histone::RFP so that transcribing nuclei could be identified and tracked in space and time (Figure 2C,D). Confocal live imaging of isolated ovaries revealed bright puncta in follicle cell nuclei corresponding to nascent transcription of the endogenous genes (Figure 2C,D; Movies S1,S2). In both cases, transcription was highly asynchronous and stochastic between nuclei (Figure 2E) as suggested by the distribution of ATS and despite Delta expression in all neighbouring cells and the requirement for Notch activity throughout the FE ^31,32,35^.

As illustrated by the kymographs (Figure 2E), *E(spl)m7-MS2* and *E(spl)m*β−*MS2* transcription profiles were characterised by active periods of several minutes, during which transcription foci could be detected, followed by periods of inactivity (Figure 2E,F). In agreement with the results from smFISH, only a low percentage of nuclei were active in any 5-minute time-window and a higher proportion of nuclei were transcribing *E(spl)m*β−*MS2* than *E(spl)m7-MS2* (Figure 2G). The proportions are approximately half those detected by smFISH, a result that is consistent with only one of the two alleles being active at any time (since only one allele has MS2 loops). The two genes had active periods of similar durations, with those of *E(spl)m*β*-MS2* on average slightly shorter (Figure 2H) and most underwent a single period of activity within the circa 1hr timeframe of imaging. For the few nuclei with two active periods (e.g. Figure 2E) the intervening periods of inactivity were highly variable (Supplementary Figure 2D). Altogether, it is evident that the transcription profiles are highly dynamic with short periods of activity occurring with relatively low frequency. However, once nuclei initiate transcription, maximal levels are achieved very rapidly (Figure 2I), with *E(spl)m*β *-MS2* reaching higher levels of mRNA production than *E(spl)m7-MS2 (*Figure 2I*). B*y calibrating accurately the MS2/MCP fluorescence against the measurements using smFISH (Supplementary Figure S2B, C and E), we estimate that a maximum of 35 RNAs are actively being transcribed at any time point during the height of *E(spl)m*β activity, with a detection threshold of 3 molecules.

Gene activity periods are usually comprised of multiple ON and OFF cycles known as transcriptional bursts ^45,46^. Consistent with this model, transcription levels of both genes fluctuated within each active period (Figure 2E,F) and we therefore used a statistical inference method, BurstDECONV, to deconvolve the MS2/MCP signal traces into individual transcription initiation events ^47,48^ (Figure 2J; Supplementary Figure 2F). This generates a distribution of waiting times between successive polymerase initiation events and the number of exponentials required to fit these data then determines the number of promoter states in the model. As the resulting distributions for *E(spl)m*β and *E(spl)m7* active periods could not be fitted by a bi-exponential model (Supplementary Figure 2G) but could be fitted with 3-exponentials, our analysis suggests the existence of 3 promoter states (Figure 2J; Supplementary Figure 2G). These consist of a competent promoter-ON state and two distinct promoter-OFF states, one of which may correspond to a polymerase paused state ^47^. The mean durations and probabilities of the 3 states were broadly similar for the two genes (Supplementary Figure 2I) and suggest that, within each active period, their promoters are rapidly switching between ON and OFF states with similar likelihoods.

In the competent, promoter-ON, state there is a variable probability of transcription being initiated, defined as kini. The main difference between *E(spl)m*β *and E(spl)m7* transcription kinetics was the magnitude of kini, which was significantly larger for *E(spl)m*β, consistent with the greater amplitude of its transcription intensity (Figure 2I,J, Supplementary Figure 2H). Thus, the two genes differ in the probability of PolII release once a promoter is in the ON state within each period of activity, as well as in the probability that a nucleus will enter into an active period.

In summary, these results demonstrate that the transcriptional response to Notch activity in the epithelium is probabilistic. Nuclei transcribe for short periods and then enter a variable inactive period before reinitiating. This pattern is not simple to reconcile with the ensemble averaging for Mam enrichment at this stage and was unexpected in a tissue where the cells are exposed to ligand over a prolonged period. Notably, it differs from the highly penetrant and synchronised response in the mesectoderm ^49^.

### Mam hubs are dynamic and precede transcription initiation

Having observed that Mam is enriched at *E(spl)-C* (Figure 1E, Figure 3A) and that the transcriptional profiles are highly dynamic, we sought to investigate the relationship between the two. To achieve this, we performed live imaging in 3 channels, combining the ParB/Int labeled *E(spl)-C* locus with Halo::Mam and with *E(spl)m7-MS2/MCP* or *E(spl)m*β-MS2/MCP (Figure 3B, Supplementary Figure 3A, Supplementary movies 3,4). The levels of Mam at *E(spl)-C* were tracked for up to 3 minutes before transcription was first detected and throughout the active period. Aligning these Mam intensity levels with the onset of MCP/MS2 transcription, revealed that there was a peak of Mam enrichment circa 2 minutes prior to transcription onset, whichever gene was tracked (Figure 3C-D).

**Figure 3:**
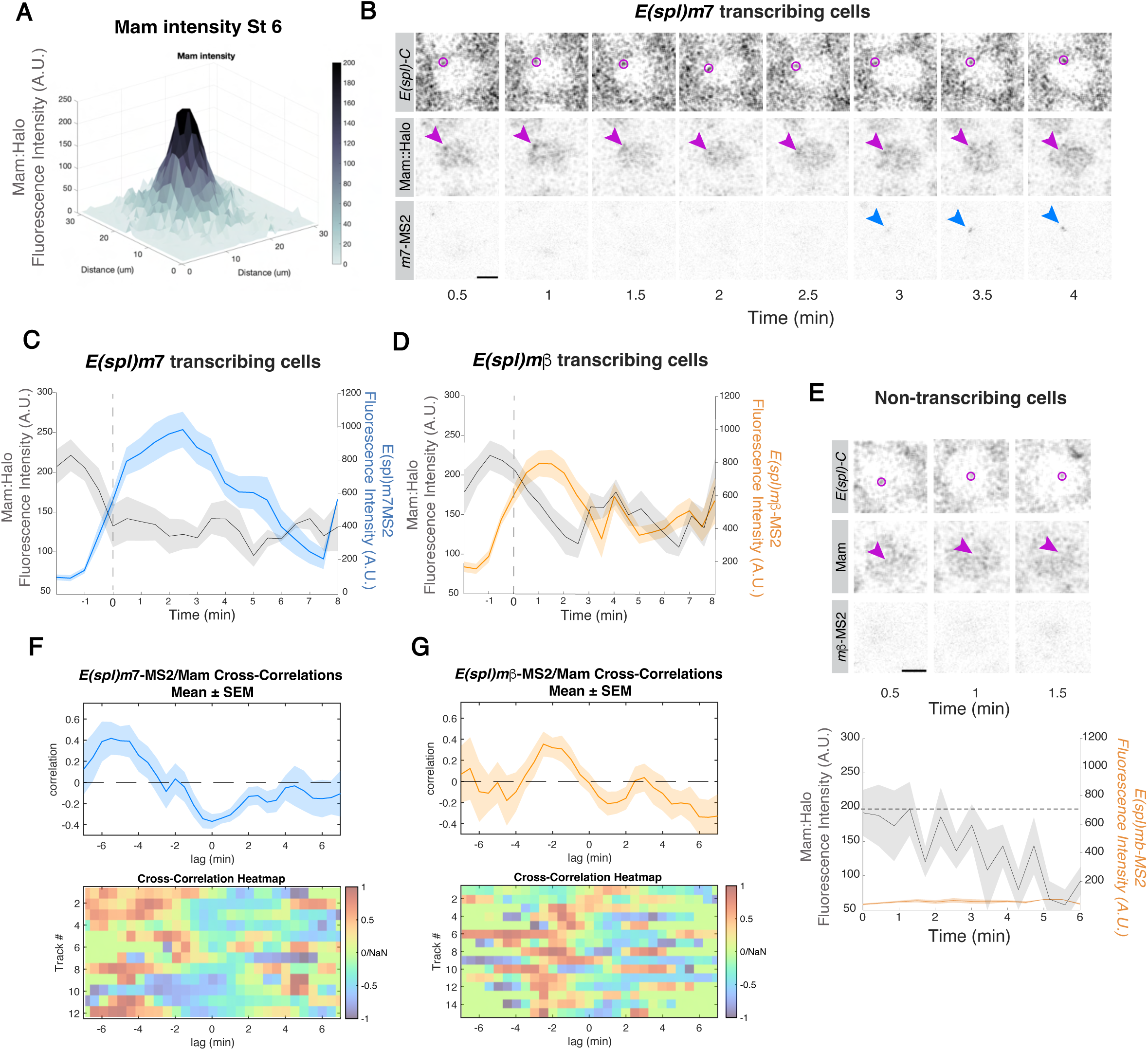
Mam hubs are dynamic and prefigure transcription. (A) Mean Mam intensity at *E(spl)-C* in stage 6, visualized by 3D surface plot. n= 141 nuclei from 3 e.c (B) Confocal images from *in vivo* movie, tracking Mam-Halo and *E(spl)m7*-MS2 in relation to *E(spl)-C* locus (magenta circle). Mam enrichment (arrowhead) precedes *E(spl)m7*-MS2 transcription (blue arrowhead). Scale bar represents 2 μm. Supplementary movie S3 (C-D) Average intensities of Mam (grey) and *E(spl)m7*-MS2 (C, blue) or *E(spl)m*β-MS2 (D, orange). Intensities are time aligned respect to transcription initiation (dotted line). *E(spl)m7-*MS2, n = 12 nuclei from 7 e.c., *E(spl)m*β-MS2, n=15 from 5 e.c (E) Confocal images from *in vivo* movie of non-transcribing nuclei as in B; no Mam enrichment occurs at *E(spl)-C.* Graph, average intensities of Mam (grey) and *E(spl)m*β-MS2 (orange) in non-transcribing cells. Scale bars represent 2 μm. n = 7 nuclei from 4 e.c (F-G) Cross-correlation analysis to test the relationship between transcription profiles and Mam enrichment. Mean correlation R^2^ value (top panels) and heatmaps (bottom panels) comparing the Mam profiles at different time lags with E(spl)*m7*-MS2 (F) and E(spl)*m*β-MS2 (G). In C,D,F,G shading represents standard error of the mean (SEM).

Mam enrichment at *E(spl)-C* was evident in all nuclei where transcription was initiated but did not reach levels above the nuclear average in nuclei that were not actively transcribing *E(spl)-C* genes, suggesting a threshold mechanism might operate (Figure 3E). These results support the hypothesis that formation of a Mam hub is a fundamental step in Notch-dependent activation of transcription but are difficult to reconcile with the cumulative Mam enrichment detected at this stage (Figure 3A), given the relatively low frequency of active nuclei. We therefore returned to cumulative intensities of Mam at *E(spl)-C* and ranked the enrichments in quartiles (Supplementary Figure 3B). This analysis revealed a gradation of enrichments, consistent with the dynamic fluctuations detected by live-imaging.

The dynamic changes in Mam enrichment at *E(spl)-C* raise the possibility of a quantitative relationship between Notch activation complexes, as represented by Mam, and the transcriptional output. To assess this, we performed a cross-correlation analysis between the dynamic Mam and MCP/MS2 profiles for each nucleus ^27,50,51^. This tests the degree of similarity between two signals as a function of the time lag between them. Strikingly, there was a positive correlation between the Mam hubs and the transcription profiles for both genes, indicating that their rates of change are related to one another (Figure 3F,G). The correlation was maximal when the intensity profiles were shifted relative to one another by 2-5 minutes (Figure 3F,G) which implies that the Mam hubs are maximal a few minutes prior to transcription being initiated. We also queried whether Mam enrichment levels correlated with the slope of the MS2/MCP transcription profiles, indicative of a direct relationship between Mam recruitment and PolII loading. There was a positive correlation for *E(spl)m7,* arguing that the two are quantitatively related for this gene (Supplementary Figure 3C). The lack of a similar correlation for *E(spl)m*β suggests an “all or none” response occurs once a threshold level of Mam is reached.

We have previously detected Notch dependent Mam enrichments associated with *E(spl)-C* in another tissue, the salivary glands. In those conditions an additional signal, provided by the hormone ecdysone, was required to bring about robust transcription. To investigate whether the dynamics between the Mam hub and transcription are conserved in this tissue, we imaged the *E(spl)-C* locus in Notch-ON, ecdysone-treated nuclei in combination with *E(spl)m*β-MS2/MCP at high resolution (Figure 4A). This tissue is unusual in having polytene chromosomes containing many aligned copies of the genome. When imaged at high resolution in cross-section we could detect heterogeneities within the large diffuse “nuage” of Mam recruited around *E(spl)-C* (Figure 4B, Supplementary Movie 5). The punctate high-density Mam regions, which we refer to as condensed hubs, were associated with *E(spl)m*β-MS2 signals such that most Mam condensed hubs were paired with MS2 foci and vice versa (Figure 4B, Supplementary Figure 4A, Supplementary Movie 5).

**Figure 4:**
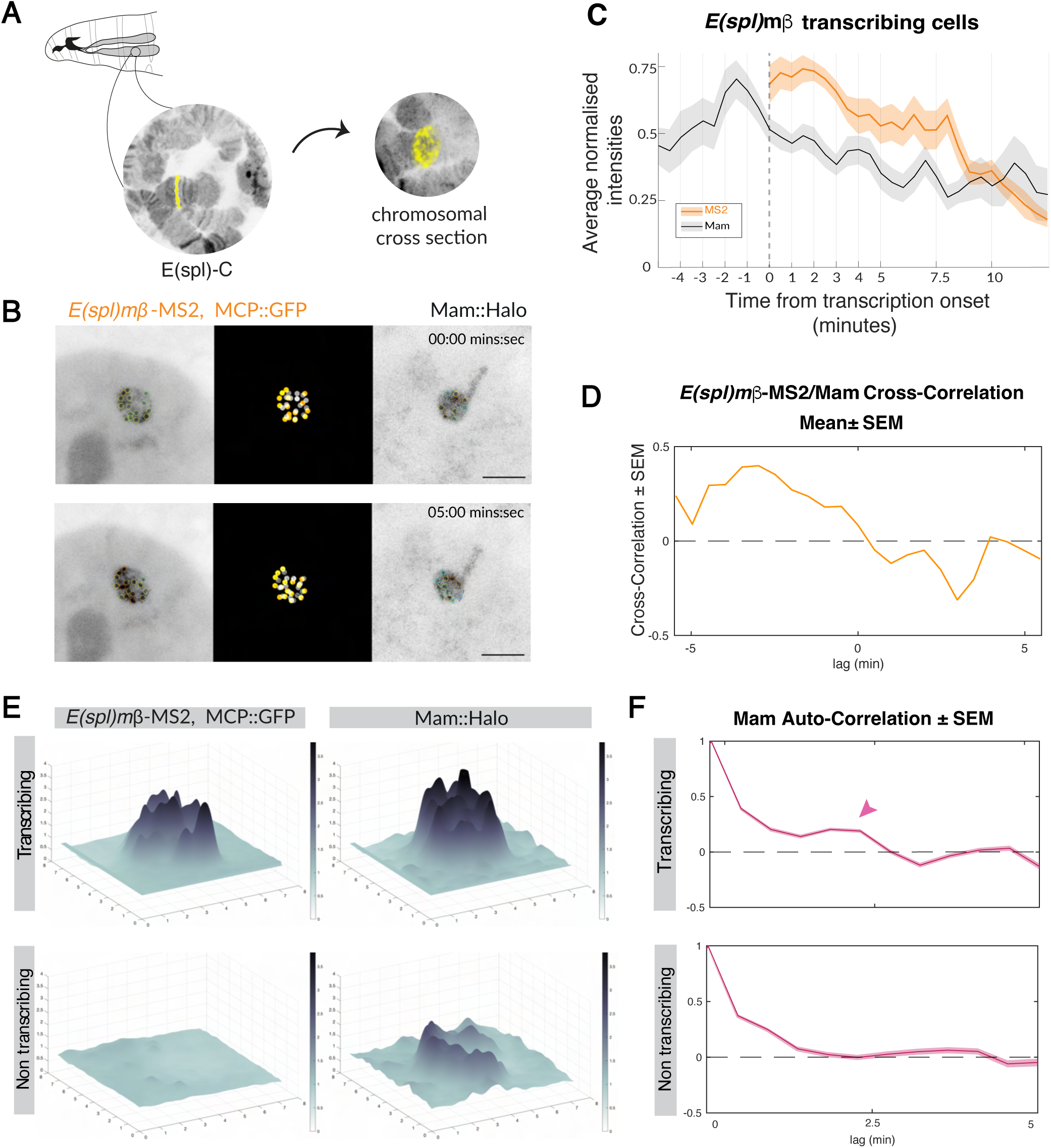
Condensed Mam hubs in salivary glands precede and correlate with transcription. (A) Cartoon illustrating imaging approach in salivary glands, with optical cross-section revealing aligned chromosomal copies as separate foci. Yellow pseudo-colouring indicates factors recruited to *E(spl)-C*. (B) Live-imaging to track *E(spl)m*β-MS2 (greyscale, left; orange, middle)) and Mam (grey, middle; greyscale, right) at *E(spl)*-C in cross-sections of salivary gland chromosomes. Pairs of dense Mam foci and *E(spl)m*β−MS2 foci can be tracked over time. Supplementary move S5. Scale bars represent 5 μm. (C) Mean intensity profiles of Mam (grey), *E(spl)m*β-MS2 (orange) aligned by transcription onset (dotted line, no quantification prior to onset), shading represents standard error of the mean (SEM). n = 20 tracks. (D) Cross-correlation analysis to test the relationship between paired *E(spl)m*β-MS2 transcription profiles and Mam enrichment profiles, mean correlation R^2^ value at different time lags. For heat maps see Supplementary Figure S4C. (E) 3D plot of *E(spl)m*β−MS2 and Mam-Halo enrichment at *E(spl)-C* in transcribing (upper panels) and non-transcribing cells (lower panels), single timepoint from Supplementary movie S6. (F) Auto-correlation analysis to test for periodicity in Mam enrichment profiles. Auto-correlation values in transcribing (upper panels) and non-transcribing cells (lower panels), reveal second peak (arrow) in transcribing nuclei.

Tracking the paired Mam and *E(spl)m*β -MS2 foci we detected a peak of maximal Mam enrichment just before transcription started (Figure 4C). Comparing quantitatively the profiles of Mam in the condensed hubs with the transcription intensity profiles, by performing a cross-correlation analysis on each pair, confirmed that the two are positively correlated (Figure 4D, Supplementary Figure 4C). As with Mam hubs in the FE, the maximum correlation was achieved when the two profiles were aligned with a lag, indicating that the Mam hub achieves maximal levels shortly before transcription is initiated and that the rates of change are related (Figure 4D). To assess Mam behaviour in nuclei that were not transcribing, we imaged tissues without ecdysone treatment. These non-transcribing nuclei had a more homogeneous diffuse “nuage” of Mam enrichment across the chromosome, as described previously, and lacked the dense foci of transcribing nuclei (Figure 4E, Supplementary Figure S4B, Supplementary movie 6; Deharo-Arbona et al., 2024). On this basis, we propose that the smaller dense condensates are analogous to those found in the FE. In both cases they are present transiently and achieve maximal levels prior to transcription. Additionally, a correlation analysis of the Mam profiles against themselves (autocorrelation; Figure 4F) revealed a small positive correlation at circa 2.5 mins suggesting there is a periodicity in the dynamics of hub condensation under these conditions of ectopic Notch activity.

Overall, the results demonstrate that Mam becomes dynamically enriched into a condensed hub that precedes and correlates with transcription being initiated at the Notch regulated genes. We propose that this is a core feature of the mechanism for initiating transcription downstream of Notch signalling.

### Mam hub is stabilised when transcription is inhibited

Mam enrichment into a local hub is related to transcriptional activity. To distinguish whether it is a cause or a consequence of transcription being initiated, we tested the consequence of pretreating the tissues with the highly specific transcription inhibitor, Triptolide, which impedes preinitiation complex (PIC) formation ^52^. Live imaging of *E(spl)m7*-MS2/MCP after tissues were treated with triptolide for one hour confirmed that transcription had been completely abolished (Figure 5B). Strikingly however, Mam levels at *E(spl)-C* were enhanced under these conditions, giving rise to a more intense and compact Mam hub (Figure 5A). A second transcription inhibitor, DRB, which interferes with the release of paused PolII also brought about a similar increase in the intensity of the Mam hubs (Supplementary Figure 5A, B). These results demonstrate that transcription per se is not required for recruitment of Mam into a hub. Indeed, they suggest the converse, that transcription/mRNA production destabilises the hub and potentially contributes to its dissolution.

**Figure 5:**
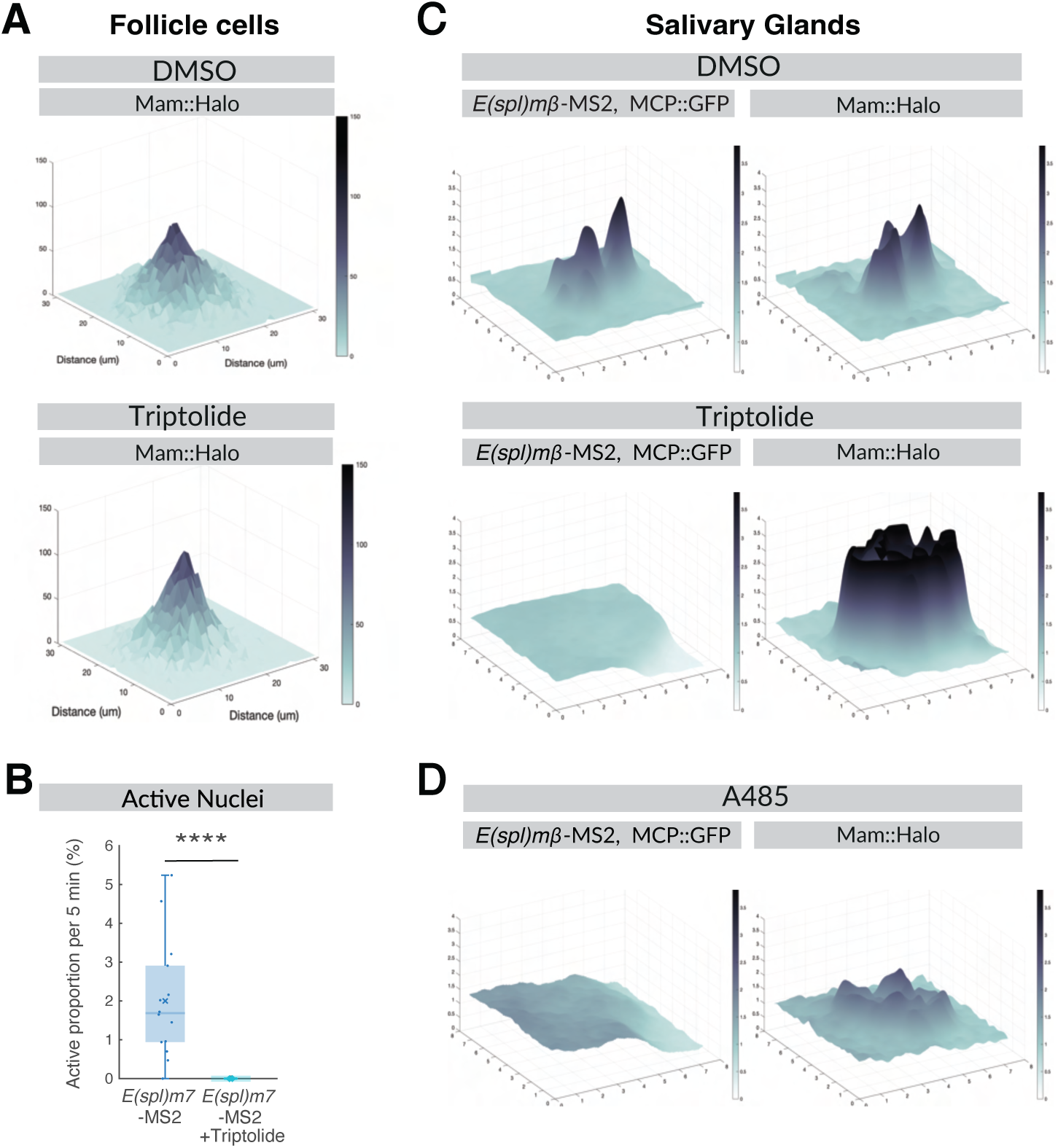
Mam hub persists when transcription is inhibited. (A-B) Effect of Triptolide treatment on Mam enrichment and transcription in follicle cells. (A) 3D surface plot of average Mam intensity levels at *E(spl)-C* in control (DMSO) or Triptolide treated tissues. n= 531, 540 from 7 and 6 e.c. (B) Average proportions of *E(spl)m7-MS2* transcribing nuclei from before and after triptolide treated (n=3) stage 6 egg chambers. (C) Effect of Triptolide treatment on Mam enrichment and transcription in salivary glands. 3D surface plot of *E(spl)m*β-MS2/MCP and Mam intensity levels at *E(spl)-C* locus in control (DMSO) or Triptolide treated tissues, representative time point. (D) Effect of A485 treatment on Mam enrichment and transcription in salivary glands. 3D surface plot of *E(spl)m*β-MS2/MCP and Mam intensity levels at *E(spl)-C* locus in control (DMSO) or A485 treated tissues, representative time point.

Comparable results were obtained in the salivary glands. Although there was little change in the diffuse Mam recruitment in non-transcribing nuclei following triptolide treatment^15^, we detected a significant increase in Mam intensities within condensed hubs of transcriptionally active nuclei, as in the FE (Figure 5C, Supplementary Movie 7). As Mam is reported to interact with the histone acetylase CBP/p300 ^13,14,53^ we also analysed the consequences of exposing tissues to A485, a potent inhibitor of its acetylase activity. CBP/p300 inhibition resulted in a significant decrease in the fluorescence intensity of the condensed Mam hubs (Figure 5D, Supplementary movie 8). The A485 treatment did not fully eliminate the dense puncta and the surrounding “nuage” was not altered (Figure 5D; ^15^, suggesting that other factors also contribute to robust Mam recruitment. However, these data reveal that Mam condensation is slowed or less efficient in the presence of A485, implying that CBP/p300 normally plays an important role similar to its positive effects on Sox2 recruitment ^54–56^.

Together these data argue firstly that the formation of condensed Mam hubs at *E(spl)-C* is independent of transcription and is enhanced by the activity of CBP/p300. Secondly, it appears that the Mam foci are normally destabilised when transcription progresses. This would generate a cyclical loop of Mam condensation and Mam dispersal in response to signalling ^57^.

### Mam hub is sensitive to Notch activity levels

Mam is a core component of the activation complex, recruited by NICD. We therefore hypothesised that the intensity of Mam hubs would be directed by the levels of Notch activity. To address this question, we used genetic strategies to increase or reduce Notch and monitored the consequences on Mam recruitment and transcription during stage 6.

To investigate the consequences from increasing Notch activity, a constitutively active form of Notch, NotchΔECD, was expressed at moderate levels throughout the tissue. Under these conditions, Mam enrichment levels at *E(spl)-C* were strongly elevated giving rise to a dense hub in many nuclei and less diffuse nucleoplasmic protein (Figure 6A,B). The effects of elevated Notch on transcription were then analysed by smFISH and by tracking transcription live with MS2/MCP. In the latter case genetic constraints resulted in conditions with a lower level of NotchΔECD being present (see methods). Two main consequences from elevated Notch were seen. First, there was an increase in the proportion of actively transcribing nuclei (Figure 6C,D; Supplementary Figure 6A). This was most evident for *E(spl)m7,* which showed a marked increase using both approaches, while for *E(spl)m*β the difference was only evident with in higher overexpression conditions, seen by smFISH, where the proportion and size of the ATS were both increased (Figure 6C,D). Second, the amplitude of transcription, measured by *E(spl)m7-MS2* and *E(spl)m*β*-MS2* transcription profile intensities, was increased for both genes (Supplementary Figure 6D) while the active and inactive period durations were unaffected (Supplementary Figure 6B,C). Thus, the elevated Notch activity results in a greater probability of nuclei entering an active phase and higher level of mRNA production within each active period, which correlates with the increase in Mam hubs.

**Fig. 6.**
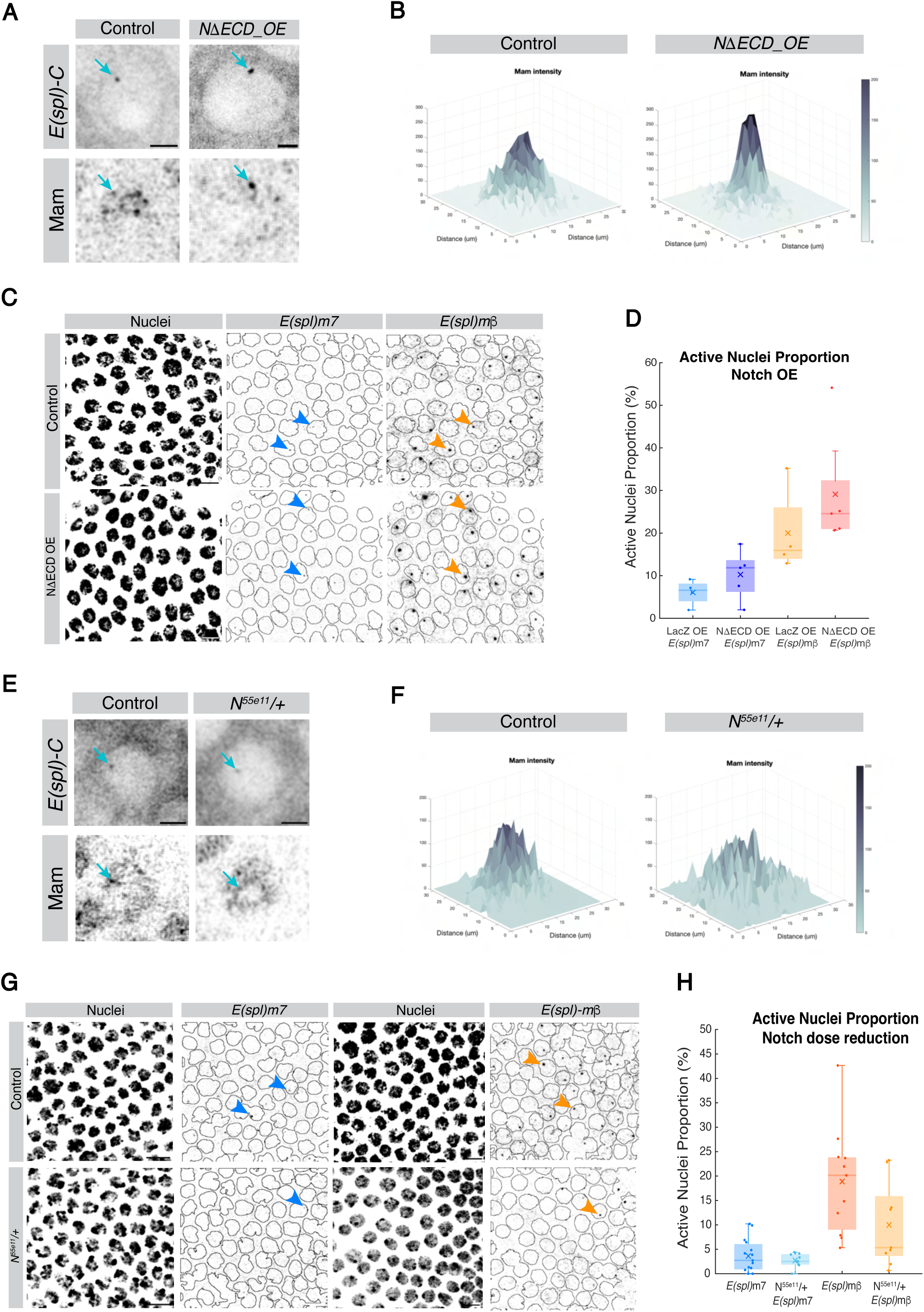
Effects of Notch activity levels on Mam enrichment and transcription profiles. (A-D) Elevated Notch activity. (A) Live imaging of *E(spl)-C* (upper panels, cyan arrows) and Mam hubs (lower panels, cyan arrows) in control (*UAS-LacZ*) (left) and Notch overexpression conditions (*UAS-N*Δ*ECD*) (right) (lower panels, cyan arrows). Scale bars represent 2μm. (B) 3D surface plot of mean Mam intensity at *E(spl)-C* in control (left) and NΔECD overexpression (right) conditions. n= 240 (*UAS-LacZ*) and n= 279 (*UAS-N*Δ*ECD*). (C) Confocal images of smFISH with *E(spl)m7-570* and *E(spl)m*β-670 probes in control (*UAS-LacZ*; upper panels) or increased *Notch* (*UAS-N*Δ*ECD*; lower panels). DAPI labels nuclei (left panels) generating masks to score ATS (right panels). Scale bars=10 μm. (D) Proportions of active nuclei in control and NΔECD detected by smFISH for both target genes. Mean data for egg chambers, n=4-6. (E-H) Reduced Notch activity. (E) Live imaging of *E(spl)-C* (upper panels, cyan arrows) and Mam hubs (lower panels, cyan arrows) in control (*UAS-LacZ*) (left) and Notch heterozygous conditions (*N^55e11^/+*) (right) (lower panels, cyan arrows). Scale bars represent 2μm. (G) 3D surface plots of Mam mean intensity at *E(spl)-C* in Control (left) and Notch heterozygous (right) conditions. n= 137 (Control) and 76 (*N^55e11^/+*). (H) Confocal images of smFISH with *E(spl)m7*-570 and *E(spl)m*β -670 probes in control (upper) or reduced Notch (*N^55e11^/+; lower panels*). DAPI labels nuclei used to form mask outlines. Scale bars = 10 μm. (I) Proportions of active nuclei in control and *N^55e11^/+*detected by smFISH for both target genes. Mean data for egg chambers, n=9-15.

To reduce Notch levels, we generated tissues that were heterozygous for a loss-of-function Notch allele (*N^55e11^*). Under these conditions, enrichment of Mam at *E(spl)-C* was decreased (Figure 6E,F) and the proportion of nuclei actively transcribing E(spl)*m7* and E(spl)*m*β, based on the ATS analysis, was reduced, (Figure 6G,H). Likewise, a lower fraction of nuclei was transcribing E(spl)*m7-MS2 (*Supplementary Figure 6E). The proportions transcribing *E(spl)m*β*-MS2* were highly variable, with no consistent reduction, suggesting the residual Notch is close to the threshold required for its activation (Supplementary Figure 6E). The amplitude of transcription during the active period, and the active/inactive period durations were unaffected (Supplementary Figure 6F,G,H).

Together these observations demonstrate that Mam recruitment is sensitive to altered Notch activity levels in a manner that allies with the consequences on gene transcription, primarily with the probability of a nucleus actively transcribing and to a lesser extent with the levels of mRNA then produced.

## DISCUSSION

Transcription factor hubs or condensates have been observed in several different biological contexts and organisms, and there is growing evidence of a functional link with transcriptional regulation ^25,27,58^. But their role in endogenous cellular processes remains enigmatic because of the challenge from probing their existence in physiological contexts. Here, by live imaging the co-activator Mam in tissues with endogenous Notch signalling, we demonstrate that high density Mam enrichments, which we refer to as condensed hubs, are induced in conditions with active signalling and that their formation precedes and correlates with gene activity, specifically with the probability and levels of transcription (Figure 7). These mature transcription-competent hubs are the final step linking Mam recruitment to promoter activation and their stepwise assembly may be a common feature of transcription factor condensates ^15,59^. Understanding how cells decode signals and generate accurate transcription outcomes is of fundamental importance for development, adult homeostasis and disease. Our data implicating a key role for transcription hubs in implementing the Notch signal sheds new light on how activity levels inform the amount of mRNA transcribed from a target gene.

**Figure. 7.**
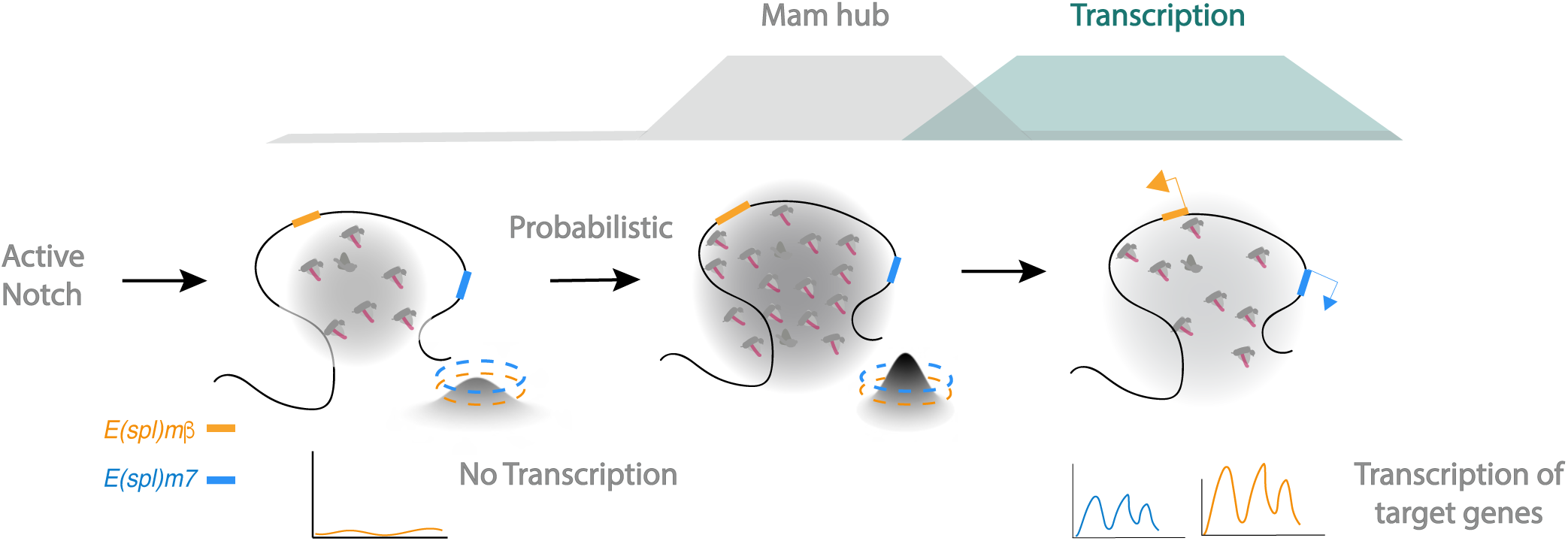
Model illustrating dynamic relationship between Mam hub condensation and transcription. Mam-containing transcription complexes (magenta symbols) assemble in response to Notch activation and are recruited to E(spl)-C, containing *E(spl)m7* (blue) and *E(spl)m*b (orange). (Left) Low levels of Mam recruitment in most nuclei (grey shading and 3D plot) are below the threshold required to promote transcription of *E(spl)m7* (3D plot, blue dash) or *E(spl)m*b (3D plot, orange dash). In a proportion of nuclei (middle), a more concentrated Mam hub is formed (grey shading) due to higher Notch levels and is sufficient to promote transcription (green shading) of one or both genes, according to their different thresholds (3D plot, orange/blue dash). Condensation of Mam precedes and correlates with transcription (top curves). Once transcription has initiated, the density of Mam at E(spl)-C is dispersed (Right) and the cycle can be reinitiated.

The characteristics of the Mam enriched condensed hubs are similar to those observed for synthetic (Gal4-VP16) and endogenous transcription factors in early *Drosophila* embryos ^25,27^ and in other contexts ^60–62^. As in those examples, localised Mam enrichments of less than 500 nm are present shortly before transcription onset and decrease in intensity as transcription progresses. Visualising the interplay between hubs and nascent transcription in single cells at high spatiotemporal resolution made it possible to uncover this relationship, which would be masked by ensemble averaging. The correlations between the hub intensities and the active transcription periods suggest that high density Mam hubs directly coordinate transcription initiation, likely through recruitment or release of RNA Pol II ^15,60,63^. The subsequent decrease in hub intensity may be a consequence of nascent RNA production, which has been shown to destabilize transcription factor condensates ^64^. In agreement, the increased intensity/stability of Mam hubs we detect with triptolide or DRB treatment argues that their dynamics are regulated by the transcription cycle, as observed for Mediator condensates ^65,66^. As degradation of NICD has also been linked to the transcription cycle this may also contribute to the dynamics of Mam recruitment ^13,67^. In addition, CBP/p300 acetylase activity appears to promote the maturation into dense Mam hubs, similar to its’ roles in the assembly of other transcription hubs^58,68^. Whichever the mechanisms, the temporal properties of Mam hub formation and dissolution provide a vehicle for transcriptional decoding of dynamic signalling inputs ^57^.

Since Mam requires NICD for assembly into the transcription complex with CSL we hypothesized that dynamic changes in Notch signalling would translate into Mam levels within the transcription hub. In agreement, locus-specific Mam enrichment was detected during stages when transcription of Notch responsive targets such as *E(spl)m7* and *E(spl)m*β occurs and was correlated with the ensuing profiles of transcription. Furthermore, changes in levels of Notch activity altered the amount of Mam within the hub in a manner that related to the probability and amplitude of transcription response. In the multi-copy *Drosophila* polytene chromosomes, where we previously detected a diffuse Notch-signalling induced hub in the absence of transcription ^15^, a change in the local distribution of Mam heralds the transition to transcriptionally active nuclei when it becomes concentrated into condensed foci that prefigure and cross-correlate with sites of active transcription in a similar manner. All of these characteristics support the model that activator-coactivator hubs form in response to signalling and confer a probability of transcription that relates to their intensity and durations (Figure 7).

As endogenous Notch signalling is maintained over a prolonged period in the follicular epithelium, we were initially surprised to find that transcription of *E(spl)m*β and *E(spl)m7* occurred sporadically. Despite this, their ATS coincide, demonstrating that firing is not random and suggesting that their probability of transcribing is related to fluctuating levels of Notch activity, as seen for Notch regulated genes in *C.elegans* gonads ^69^. In agreement, both *E(spl)m*β and *E(spl)m7* had a higher probability of transcribing when the Notch activity levels were mildly increased. Of the two genes, *E(spl)m7* had a significantly lower probability and amplitude of transcription, a pattern that is also evident in single cell RNA data from this tissue ^41,42^ suggesting *E(spl)m7* requires a higher Notch threshold than *E(spl)m*β (Figure 7) Concordantly*, E(spl)m7* transcription profiles were more sensitive to changes in Notch activity.

The differing profiles also imply that a Notch induced Mam hub of a given size/density yields different outcomes depending on the promoter affected. In a similar manner, different target genes of the Dorsal transcription factor showed distinct requirements with respect to the properties of the Dorsal hub ^27^. Explanations likely lie in sequences present in the enhancers/promoters that will recruit additional factors. For example, these might prime *E(spl)m*β so that it is poised for activation or repress *E(spl)m7*, to confer a more refractory state ^49,70^. As a result, one gene is more frequently activated than another under conditions where Notch activity levels are close to the threshold. This type of probabilistic behaviour may be common not only for Notch regulated genes but also much more widely in signalling induced responses as many genes have demonstrated distinct discontinuous transcriptional activities that result in high degrees of cellular heterogeneities.

## Materials & Methods

### Experimental animals

Species: *Drosophila melanogaster*. Flies were grown and maintained on food consisting of the following ingredients: glucose 76 g/l, cornmeal flour 69 g/l, yeast 15 g/l, agar 4.5 g/l, and methylparaben 2.5 ml/l.

### Fly stocks

Full genotypes are summarised in Supplementary Table S1. To image *E(spl)-C* locus, a recombinant “locus tag” chromosome ^16^ consisting of *Int1* sequences inserted into *E(spl)-C* intergenic region adjacent to *E(spl)mdelta* (Chromosome 3R 26038865:26038884) and *UAS-ParB1-mcherry* or *UAS-ParB1-GFP* inserted *AttP.86Fb* was used in combination with *traffic-jam-Gal4* to drive ParB1 expression in the FE ^71^ or with *1151-Gal4* to drive expression in salivary glands ^16^. To analyse expression and enrichment of TFs and other proteins we used endogenously tagged fluorescent labelled proteins generated by CRISPR, Halo::Mam, GFP::Mam ^15^, Sox14::GFP (BL-55842), Dl::mScarlet ^72^ or by genomic rescue constructs Halo::CSL ^15^, GFP::CSL, GFP::Hairless^16^.

For MCP/MS2 live imaging of transcription, 24MS2 loops and the lacZ coding sequences were inserted into *E(spl*)*m7-HLH* and *E(spl*)*mβ-HLH* genes by CRISPR-Cas9 genetic engineering. In *E(spl*)*m7*-MS2 the insertion sire was close to the C-terminus of the coding sequences using the strategy described ^49^ and in *E(spl*)*mβ*-MS2 the insertion site was closer to the transcription start-site ^15,72^. The MS2 tagged genes were combined with *hsp83-MCP::GFP* (BL-7280) to track transcription and His2Av-RFP (BDSC #23650) to label nuclei. *hsp83-MCP::GFP* (BL-7280) was also recombined with traffic-jam(tj)-Gal4 to generate a strain for triple labelling experiments with Halo::Mam, *E(spl)-C “*locus tag” and the *E(spl*)*m7*-MS2 or *E(spl*)*mβ*-MS2 chromosome. For similar experiments in the larval salivary glands, flies were generated containing *1151-Gal4*, *hsp83-MCP::GFP*, *E(spl)m*β -MS2 and Halo::Mam, *UAS-NΔECD*.

Notch overexpression was achieved either by combining tj-Gal4::tub-Gal80^ts^ with *UAS-NΔECD* ^15^, analogous stocks with *UAS-LacZ* were generated as a control, or, in experiments with MS2/MCP by combining the same UAS lines with a “leaky” Flip-out cassette (Tub>FRT.STOP.FRT>Gal4::UAS-TandemTomato). Crosses were maintained at 18°C (Notch at endogenous levels) or 29°C degrees for 24 hours (Notch overexpression). For Notch dose reduction, N^55e11^FRT19A was combined with the relevant Mam:Halo and MS2/MCP and yw was used as control.

## Method details

### Tissue preparation, drug treatments and live imaging conditions

Ovaries were dissected and mounted in Schneider’s medium (Sigma/Merck S0146) supplemented with 15% (v/v) foetal bovine serum (Sigma-Aldrich, F2442), 0.6% (v/v) streptomycin/penicillin antibiotic mix (Invitrogen 15140-122), and 0.20 mg/ml insulin (Sigma 15500). For imaging, ovaries were mounted into 35-mm poly-D-lysine-coated glass bottom dishes (Mattek, P35GC-1.5-10-C). For drug treatments, dissected ovaries were incubated with Compound E (4 hours, 100nM; Abcam HY-14176), DRB (1 hour, 500μM; Santa Cruz Biotechnology, 53-85-0) or Triptolide (1 hour, 10 µM; Sigma-Aldrich T3652) prior to imaging. For experiments testing whether transcription was abolished, triptolide was added at 10 µM after imaging was initiated. Preparation of salivary glands was as described previously^15,16^.

Live confocal fluorescence imaging of ovaries was performed on a Leica SP8 microscope equipped with seven laser lines (405, 458, 488, 496, 514, 561, and 633) and using a 40x apochromatic 1.3 oil immersion objective and two hybrid GaAsP detectors. For Mam enrichment assays Individual egg chambers were imaged with a 4x zoom, 1024×1024 pixel resolution, pinhole set to 1.5-Airy, three line-averages, a 12-bit depth, and 400 Hz scanning speed. Z-stacks were 0.5 µm wide and a total of circa 30 stacks were acquired to cover most of the epithelium at the imaged surface of the egg chamber. For MS2/MCP movies, individual egg chambers were imaged at 3.5 zoom, 512×512 pixel resolution, pinhole set to 3.5-Airy, 8-bit depth and 400 Hz scanning speed. Z-stacks of 0.5 µm width for 30 stacks were acquired. Time frame was 20 seconds. The same settings for MCP-GFP detection: 29.36 mW 488 nm argon laser detected with a hybrid detector.

For Mam enrichment together with MS2/MCP movies, egg chambers were imaged using the ×63/1.4 NA HC PL APO CS2 oil immersion objective. at 3.5 zoom, 512×512 pixel resolution, pinhole set to 1.5-Airy, 12-bit depth and 600 Hz scanning speed. Z-stacks were 0.5 µm wide and circa 15 stacks were acquired. Time frame was 30 seconds. Similar conditions were used for salivary glands, except zoom was 10x, pinhole set to 2-Airy.

### Single Molecule in Situ Hybridisation

Custom smFISH probe sets for *E(spl)m7-HLH* and *E(spl*)*mβ*-HLH genes were designed with Stellaris Probe Designer (Bioscience Technologies). Ovaries were dissected in non-supplemented Schneider’s medium and fixed for 30 min in 3.7% formaldehyde:PBS at room temperature. Ovaries were then washed three times in PBST (PBS +0.1% Triton X-100) and transfered to Methanol stepwise with 5 min incubations in PBST:MeOH 7:3, PBST:MeOH1:1, PBST:MeOH 3:7 on a nutating mixer before transferring to 100% MeOH. After 10 mins in 100% MeOH, tissues were re-hydrated with the same solutions in the reverse order (3:7 PBST:MeOH, 1:1 PBST:MeOH, 7:3 PBST:MeOH). After rinsing with PBST, ovaries were pre-hybridized with WB-A (Biosearch Technologies SMF-HB1-10), for 10 min and then transferred to hybridisation mixture prepared by adding probes to 100 µl of RNA FISH Hybridization Buffer (Biosearch Technologies SMF-WB1-60) at 1:100%v/v from 12.5 µM stock. They were then incubated at 37°C for 16-18 hours in the dark, washed twice for 30 minutes with pre-warmed WB-A, rinsed with PBST and then mounted using Vectashield with DAPI.

smFISH samples were imaged using a Leica SP8 microscope equipped with different laser lines (405, 458, 488, 496, 514, 561, 594 and 633) and a 40x apochromatic 1.3 oil immersion objective and one hybrid GaAsP detector was used. Individual egg chambers were imaged with a 4x zoom, 1024×1024 pixel resolution, pinhole set to 1-Airy, three line-averages, a 12-bit depth, and 400 Hz scanning speed. Z-stacks were 0.5 µm width and a total of around 30 stacks.

## Analysis

### Image analysis pipelines

To measure enrichments, images were analysed using MATLAB by importing Leica Images with BioFormat package (MATLAB R2021b, MathWorks and openmicroscopy.org). Using a custom MATLAB app, a squared region of interest (ROI), 30 × 30 pixels centred on the *E(spl)-C* was selected (adapted from DeHaro-Arbona et al., 2024). Fluorescence levels of Mam::Halo and other factors were then measured within this ROI, averaged per experimental condition and normalized with an average value, calculated from ROIs placed at random in the same set of nuclei. Average pixel plots were generated and profile graphs, ROIs were averaged in the y-dimension, and the mean and SEM were represented for each condition.

For quantification of Mam enrichment together with MS2 movies, *E(spl)-C* was tracked with Trackmate and the resulting ROI was assigned to the Mam and MS2 channels and fluorescent intensities obtained using a custom MATLAB script. Both Mam and MS2 signals were normalized by subtracting the background level (minimum value from the trace) and median filtered to smooth the traces.

For smFISH images, nuclei were segmented in 2D using Weka Segmentation (Fiji plugin) to detect Hoechst staining and masks were generated using watershed after transformation to a binary image and their number indicated the total number of nuclei analyzed per egg chamber. Masks were applied to the fluorescent probe channels to eliminate signal coming from the cytoplasm and a minimal threshold was used to identify the prominent ATS. The Fiji plugin Analyze Particles was used to count total number of ATS in an image and totals were divided by the number of nuclear masks and represented as proportion of Active Nuclei.

For MS2/MCP movies, the His2Av-RFP or siRDNA (Spirochrome SC007) signal was used to segment and track the nuclei in 2D. Each movie was segmented using Weka Segmentation (Fiji plugin) and watershed after transformation to a binary image to obtain masks. Tracking of the nuclei was performed using Trackmate (Fiji plugin) detecting the nuclei as masks and allowing to track a maximum distance of 3 µm, a maximum gap distance allowed of 3 µm and maximum frame gap of 2 frames. A custom MATLAB script (MATLAB R2021b, Mathworks), adapted from (Falo-Sanjuan et al., 2019), was used to detect and track the transcription spots based on the 3D Gaussian method developed in^44^. The tracked transcription spots were overlapped with the ROIS (masks) of the tracked nuclei, and spots lying outside nuclei were removed from the analysis. We note that in Notch overexpression conditions, a background subtraction was necessary in the MS2 channel to a 3px radius, before tracking transcription spots, because of heterogeneous nuclear levels of MCP-GFP.

To measure Mam::Halo and *E(spl)m*β*-MS2*/MCP::GFP dynamics in the salivary gland, the intensities of each fluorophore were tracked with the TrackMate plugin in FIJI. The resulting tracks were imported into MATLAB, where the tracks were sorted according to time of onset for MS2/MCP detection and the intensities extracted. Pairs of Mam and MS2/MCP foci were then selected based on proximity, with maximum distance 1 µm. The tracks were checked for tracking errors and normalised, with 0 and 1 as the lowest and highest intensity values, respectively, due to the large variation in intensity between tracks. The normalised and aligned data were combined to produce average intensities and distances with SEM.

For the measurements of Mam fluctuations in drug treatment experiments, movies were registered in FIJI by using the Mam channel. They were imported into MATLAB to apply Gaussian filtering and bin the signal into a 10×10 grid. The regions containing more than 1.5-fold intensity in comparison to the nuclear levels were retained, as the ones with persisting intensity over 5 frames (2.5 minutes). The max levels and standard deviation of each binned region was calculated for each movie, including 15 minutes for each one.

### MS2 Data Processing and parameters calculation

Only MS2/MCP transcription foci tracked for more than 5 frames were retained and a maximum gap of 2 frames was allowed. This produced an initial segregation of active (nuclei ON for at least 5 frames) and inactive nuclei. Heatmaps were generated for each movie to display only the active nuclei through time. To avoid false negatives arising from nuclei moving out of the field of view, only traces that started after the nuclei had been segmented and finished before their track ends were kept for analysis. Short gaps during ON periods (less than 5 frames) were interpolated using the mean intensity value of the surrounding frames. The proportion of active nuclei was defined as the percentage of nuclei that come ON per 5 minutes period in a maximum of 60 minutes time window per movie. The amplitude was calculated by alignment of the onset of the ON periods and values of 0 were assigned when the trace ended in order to account for proportion of active nuclei as well as intensities.

For the calculation of active periods, the same procedure was followed, except the allowable short gaps were extended to fewer than 21 frames (approximately 7 minutes). Active periods were only calculated when found between two inactive periods. Similarly, inactive periods were quantified when found in between two active ones.

### MS2 signal calibration

To estimate the average number of mRNA molecules present in MS2 foci from *in vivo* movies, we first calculated the fluorescence of a single mRNA molecule, using a method similar to^47,73^. Briefly, probe sets with different fluorophores for the gene of interest were hybridized to the samples. Then, after applying a threshold to avoid non-specific particles, the median intensity value of all detected particles was used as a proxy for the intensity of a single particle of mRNA. Active transcription sites (ATS) total intensities were divided by the intensity obtained for a single particle to determine the number of RNA molecules being transcribed. A Q-Q plot was then used to relate the intensities of ATSs, in terms of number of molecules from smFISH experiments, to the arbitrary fluorescence intensities of MS2 foci from the same gene. This yielded a linear relationship with similar parameters for each. The data for the two genes were therefore combined into a single Q-Q plot and the parameters obtained from the linear fit were used to provide the conversion factor to determine the number of mRNA molecules transcribed in the MS2 transcription data.

### Modelling

Signal deconvolution and kinetic model inference were performed using the BurstDECONV framework ^48^. After calibration, MS2 fluorescence traces for both *E(spl)m7-MS2* and *E(spl)mβ-MS2* were segmented to isolate only the active periods (Supplementary Figure S2F), thereby excluding prolonged transcriptionally silent periods. This pre-processing step ensured comparability between the two genes, as the inactive periods *E(spl)m7-MS2*, due to their long durations, were rarely captured within the timescale of our experiments. Each MS2 trace was subsequently deconvolved to infer the timing of individual RNA Pol II initiation events using the parameter settings shown in Supplementary Figure S7A.

Using the deconvolved initiation events, BurstDECONV computed a survival function describing the distribution of waiting times between consecutive RNA Pol II initiation events. This empirical distribution was subsequently fitted using both bi-exponential and tri-exponential models:

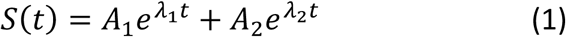

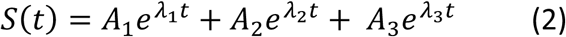

The minimum number of exponential components required to adequately fit the survival function was selected and interpreted as the number of underlying promoter states.

Model fit quality was evaluated using two complementary approaches. First, confidence intervals for the empirical survival function were calculated using Greenwood’s formula. As an initial validation step, we confirmed whether the best-fit parametric survival curve fell within these confidence bounds. For both *E(spl)m7-MS2* and *E(spl)mβ-MS2*, the two-state model failed this criterion and was therefore excluded from further consideration while the three-state model satisfied this condition for both genes (Supplementary Figure 2G). To further assess model accuracy, the Kolmogorov–Smirnov (K–S) test in combination with the mean squared deviation (used as the objective function) were employed as quantitative metrics of goodness-of-fit. Our analysis supports a non-sequential three-state model (Figure 2F) but does not exclude a sequential alternative, where the promoter transitions obligatorily through the transient OFF1 state to move between the Prom ON and the long Prom OFF2 states. However, this sequential arrangement implies a strong dependence between the two inactive states, a relationship for which a plausible mechanistic basis is lacking.

The multiexponential distribution fits were used to infer kinetic parameters for Markovian models of transcriptional bursting under steady-state conditions. The resulting kinetic estimates are provided in Supplementary Figure S2I, alongside the corresponding objective function values and Kolmogorov-Smirnov (K-S) test statistics. We note that *kini*, the parameter showing the greatest difference between *E(spl)m7* and *E(spl)mβ*, was also the most reliably inferred in method validation using synthetic datasets ^48^.

Additionally, preliminary estimates of the decay constants and population proportions (the λ and A parameters in equations (1) and (2)) were obtained using a computationally efficient and numerically stable algorithm based on solving a linear system derived from cumulative integrals (https://github.com/juangburgos/FitSumExponentials/tree/main). These estimates (Supplementary Figure S7A) were subsequently used to initialise the linear regression procedure in BurstDECONV, substantially accelerating the fitting process.

As reliable extraction of kinetic parameters required that the system had reached a steady-state regime, we computed the average interval between successive Pol II initiation events (⟨τ⟩) within a narrow, temporal window. This average interval has been shown to be proportional to the reciprocal of the product of pON (the likelihood of the promoter being in Prom ON state) and kini, the initiation rate during the active state ^74,75^. Hence stability in ⟨τ⟩ across different windows served as an indicator that the underlying kinetic parameters remained unchanged over time. A window width of 4 frames (corresponding to 80 seconds) was chosen, which provided enough initiation events to yield a robust estimate of ⟨τ⟩. To assess the precision of these estimates, we applied a bootstrap resampling approach to derive 95% confidence intervals for both genes under investigation (Supplementary Figure S7B).

## Statistical Analysis

Two-tailed *t*-test was applied in cases with n > 30 for both conditions tested. Otherwise, normality was checked via a Shapiro–Wilk test. When one or more of the samples were not normal, a Wilcoxon rank sum test was applied. When more than 3 groups were compared, ANOVA test was calculated. In all cases, significance was presented as: *p<0.05, **p<0.01, ***p<0.001, and ****p<0.0001, and p-values are provided in Figure legends.

## Data and code availability

https://gitlab.developers.cam.ac.uk/cr607/ms2_analysis/-/tree/b909366aac5a72d5b74064e03cd995ee9f12282c/

## Acknowledgements

We thank our colleagues and members of our group for discussions and helpful suggestions. We are grateful to Kat Millen for assistance with Fly injections and technical resources and to Cambridge Advanced Imaging Centre for microscopy support. This work was supported by an Investigator Award from Wellcome Trust to SJB, a Herchel Smith postdoctoral fellowship to CSM, and a PDN-Wolfson College PhD studentship to CR.

