## Supplementary figures and tables for "Direct observation of Notch signalling induced transcription hubs mediating gene-expression responses"

### Santa Cruz Mateos: Supplementary Figures, Tables and movies

**Supplementary Figures**

Figure S1. Related to Figure 1. Nuclear levels remain constant during the period of Mam enrichment at *E(spl)-C* locus which requires Notch activity.
Figure S2. Related to Figure 2. Calibration of MS2/MCP intensities and results from modelling promoter states.
Figure S3. Related to Figure 3. Variations in Mam enrichment levels
Figure S4. Related to Figure 4. Analysis pipeline for Mam enrichment and transcription profiles in salivary gland nuclei.
Figure S5. Related to Figure 5. Transcription inhibition with DRB stabilizes Mam hub
Figure S6. Related to Figure 6. Changes to transcription output when *Notch* dose is altered.
Figure S7. Related to Methods. Parameters and Steady state confirmation for modelling

**Supplementary Tables**

Table S1: Details of Drosophila lines used.
Table S2: Detailed genotypes for each Figure.
Table S3: "n" numbers and p-values from statistical tests for each Figure.

**Supplementary Movies**

Movie S1: Live imaging of E(spl)m7-MS2/MCP::GFP transcription, stage 6.
Movie S2: Live imaging of E(spl)mβ-MS2/MCP::GFP transcription, stage 6.
Movie S3: 3 colour live imaging of E(spl)-locus (yellow), Mam::Halo (magenta), E(spl)m7-MS2/MCP::GFP transcription (blue), stage 6.
Movie S4: 3 colour live imaging of E(spl)-locus (yellow), Mam::Halo (magenta), E(spl)mβ-MS2/MCP::GFP transcription (orange), stage 6.
Movie S5: Live imaging of E(spl)mβ-MS2/MCP::GFP transcription (green in overlay) and Mam::Halo (magenta in overlay), salivary gland nucleus.
Movie S6: 3D plots from live imaging of Mam::Halo and E(spl)mβ-MS2/MCP::GFP transcription in transcribing and non-transcribing salivary gland nuclei.

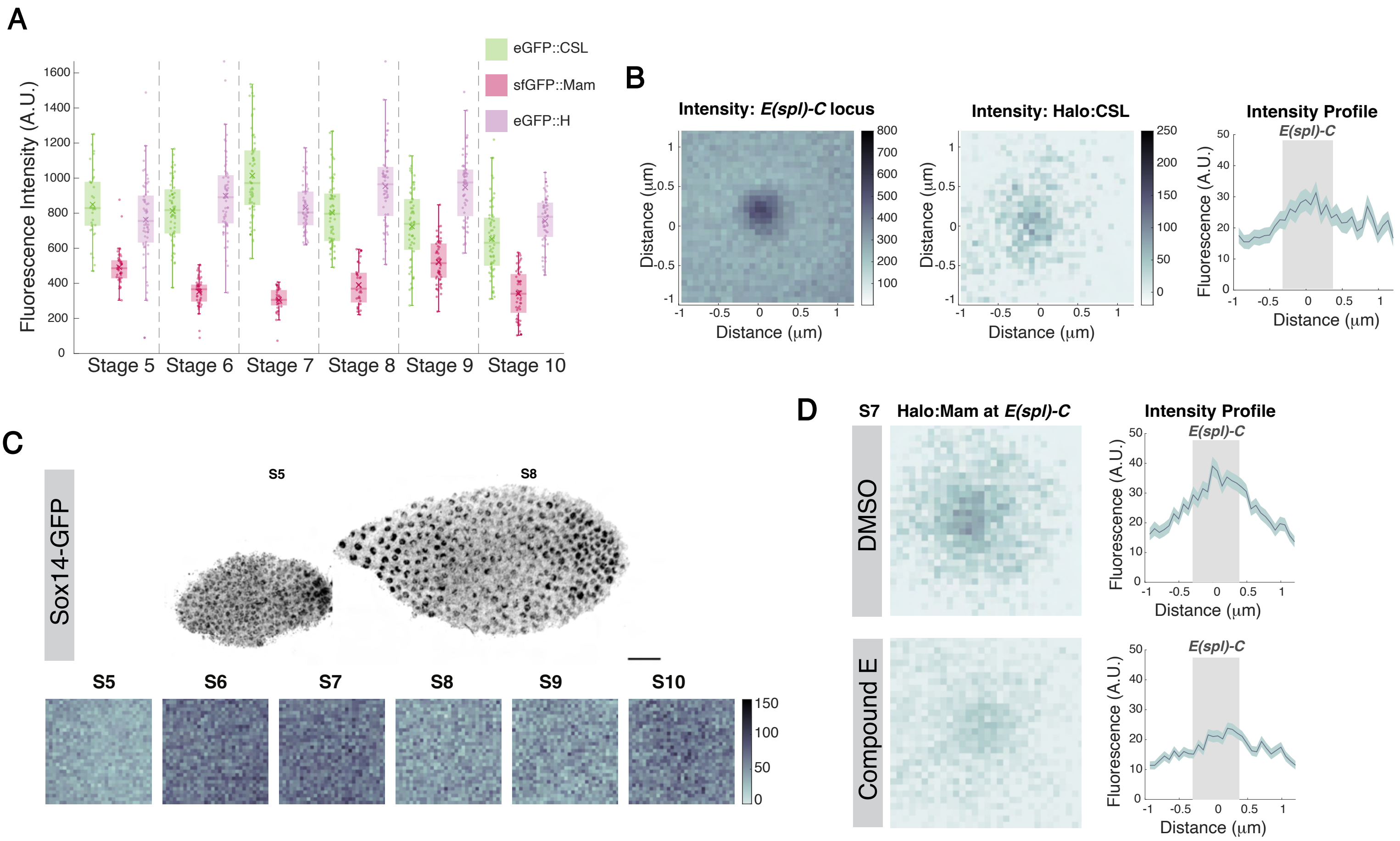


**Figure S1: Nuclear levels remain constant during the period of Mam enrichment at E(spl)-C locus which requires Notch activity.**

**(A)** Nuclear levels of eGFP:CSL, eGFP:H and sfGFP:Mam remain the same throughout stages 5-9 development. n= 36, 61, 61, 61, 61, 61 (eGFP:CSL), 61, 59, 61, 61, 61, 61 (eGFP:H) and 46, 53, 37, 37, 57 and 57 (sfGFP:Mam). **(B)** Example illustrating pipeline to quantify enrichment at *E(spl)-C* locus. Images are centred with respect to the tagged *E(spl)-C* locus (left) and the intensity values obtained from the aligned images of the fluorescent transcription factor are averaged and normalized to create an average intensity pixel map (right). **(C)** Expression of Sox14-GFP in stage 5 and 8 (upper panel). Scale bar = 20µm. Average intensity of Sox14-GFP at *E(spl)-C* during stages 5 to 10 shows no enrichment. n = 70, 123, 122, 78, 72 and 95, respectively. **(D)** Effects of Notch inhibition on Mam enrichments. Average pixel plots and intensity profiles as in A with average Mam intensities at *E(spl)-C* in control (DMSO, upper panels) and γ-secretase (Compound E) treated egg chambers (lower panels). Grey shading indicates *E(spl)-C* locus. n= 198 (DMSO), 194 (Compound E).

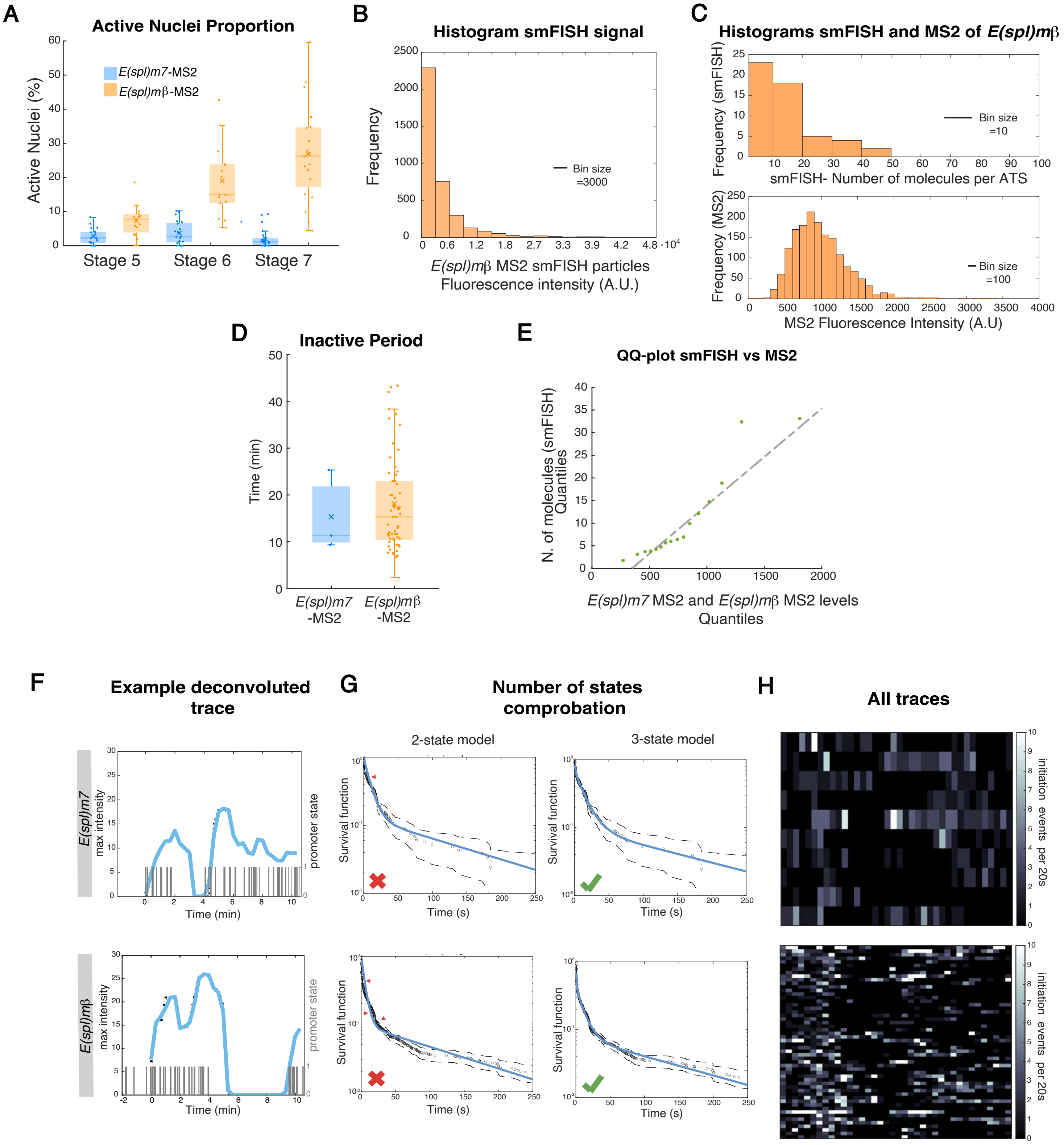


I

| | k1+ | k1- | k2+ | k2- | kini | p1 | p2 | p3 | Obj | K-S test p-value |
|---|---|---|---|---|---|---|---|---|---|---|
| ***E(spl)m7*** | 6.56e-3 | 2.94e-2 | 1.77e-1 | 4.10e-1 | **3.04e-1** | 5.75e-1 | 2.97e-1 | 1.28e-1 | 5.00e-3 | 9.99e-1 |
| ***E(spl)mβ*** | 7.02e-3 | 5.11e-2 | 1.40e-1 | 3.26e-1 | **6.52e-1** | 6.86e-1 | 2.20e-1 | 9.42e-2 | 1.06e-2 | 9.99e-1 |

**Figure S2: Calibration of MS2/MCP intensities and results from modelling promoter states.**

**(A)** Proportions of nuclei actively transcribing *E(spl)m7* (blue) and *E(spl)mβ* (orange) at the stages indicated using smFISH. **(B)** Example of calibration process. Frequency histogram of particle intensities from an smFISH image labelled with *E(spl)mβ*-670. **(C)** Histograms of ATS calibrated signals (X-axis is number of molecules, upper graph) of smFISH image of *E(spl)mβ*-670 and intensities of *E(spl)mβ*-MS2 transcription foci (lower graph). **(D)** Time between two active periods from *E(spl)m7*-MS2 (blue) and *E(spl)mβ*-MS2 (orange) traces containing >1 active period. **(E)** Combined Q-Q plot of number of molecules in ATS from smFISH images with respect to MS2 intensities from a representative movie. **(F)** Examples of results from Burstdeconv, with transcription profile (blue) and inferred PolII events (black lines).**(G)** Graphs testing fit to 2-state or

3-state promoter models. Red arrowheads point to parametric survival curve not fitting in the confidence intervals. **(H)**Heat map of PolII initiation events assigned to transcription tracks from *E(spl)m7*-MS2 and *E(spl)m*β-MS2. **(I)**Table showing the resultant values from the modelling. The biggest differences found between *E(spl)m7*-MS2 and *E(spl)m*β-MS2 is **kini**.

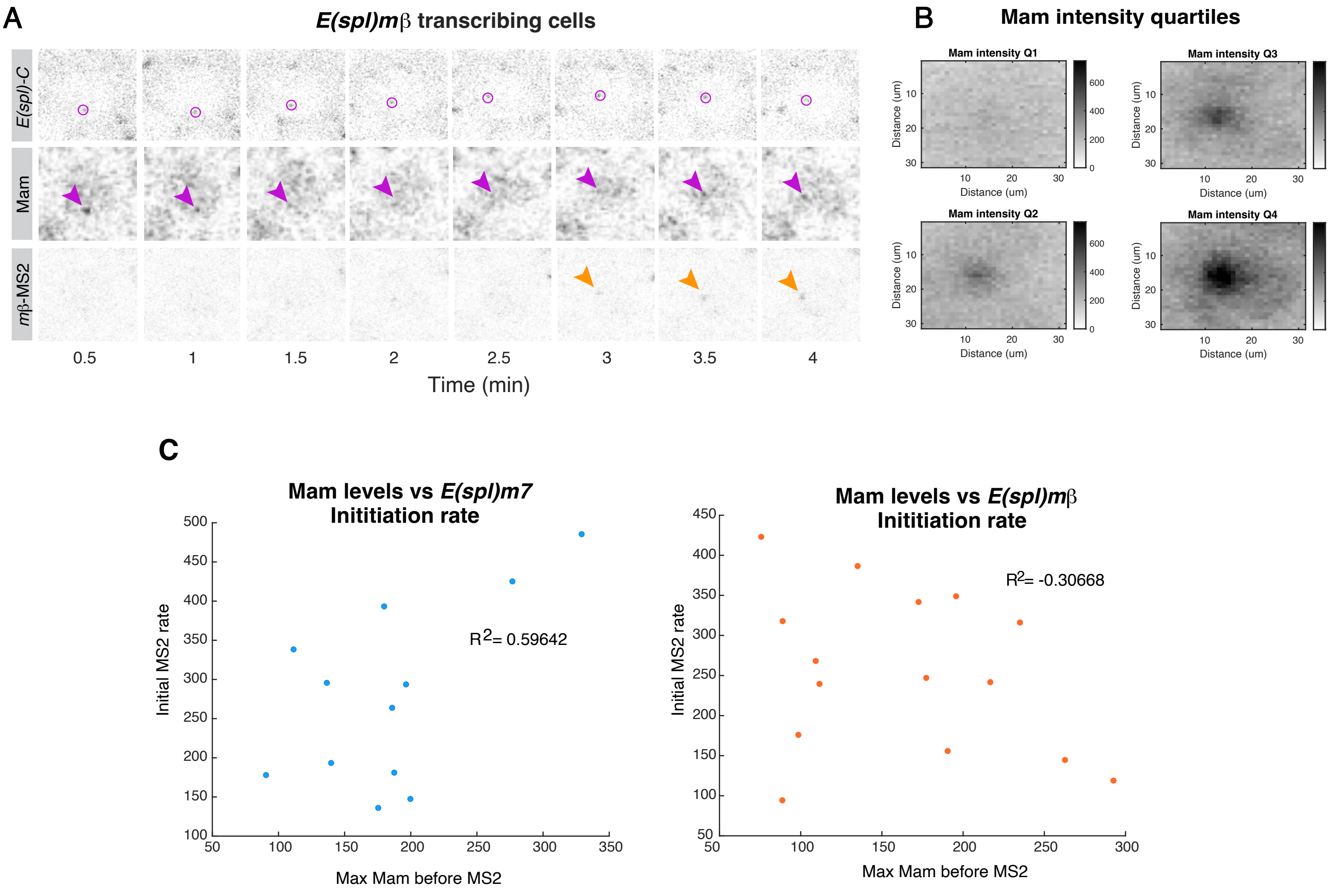


**Figure. S3. Variations in Mam enrichment levels.**

**(A)** Confocal images from *in vivo* movie (Supplementary movie S4), tracking Mam-Halo and *E(spl)m*β-MS2 in relation to *E(spl)-C* locus (magenta circle). Mam enrichment (arrowhead) precedes *E(spl)m*β-MS2 transcription (orange arrowhead). Scale bar represents 2 $\mu$m. **(B)** Average pixel intensities of Mam enrichment at *E(spl)-C* in St 6 nuclei from Figure 1D,3B have been partitioned into quartiles which reveals variations in the levels of enrichment (n= 141). **(C)** Correlation between maximal Mam intensity values and PolII loading rate, inferred from the initial slope of *E(spl)m7*-MS2 transcription profile (R2 = 0.596) and *E(spl)m*β-MS2 (R2=-0.306).

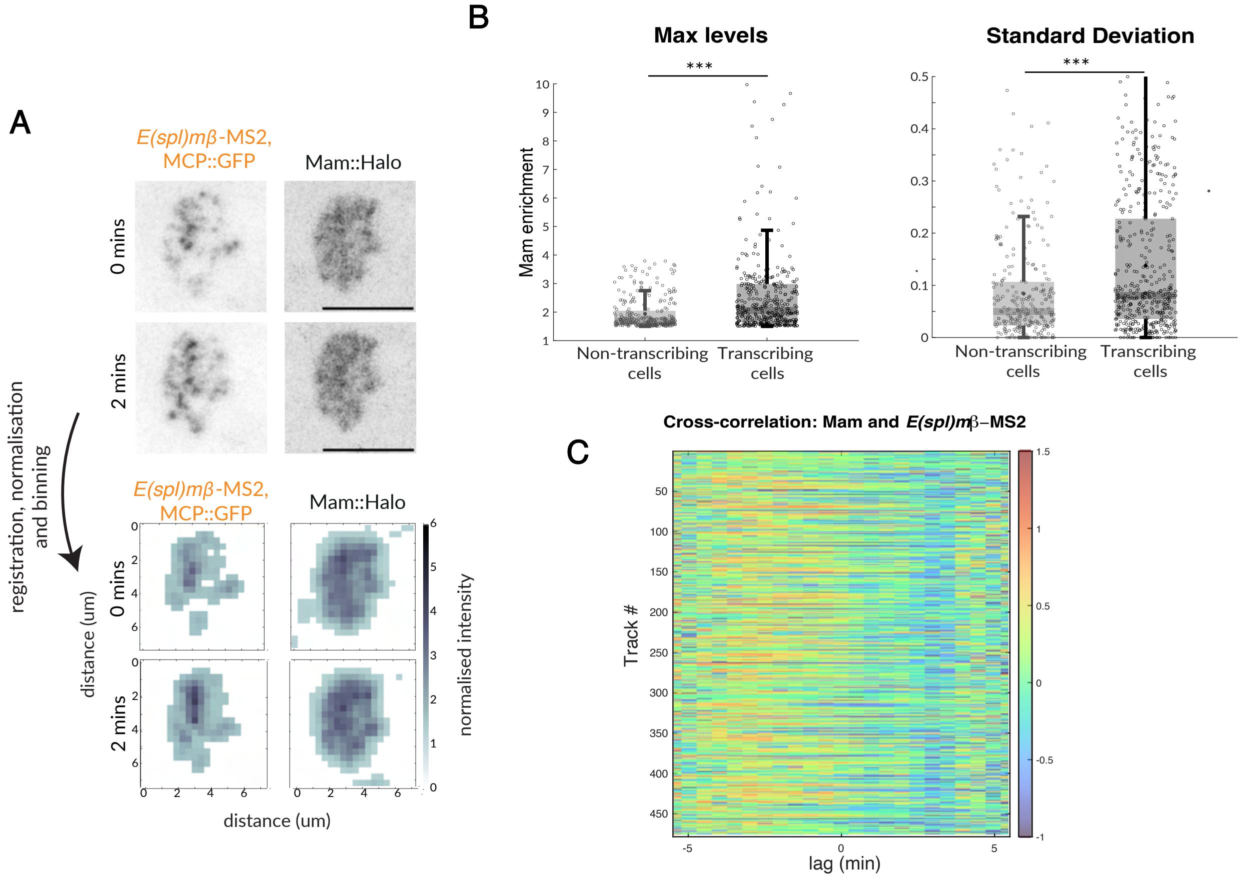


**Figure S4: Analysis pipeline for Mam enrichment and transcription profiles in salivary gland nuclei.**

**(A)** Cartoon illustrating analysis pipeline to measure intensities of *E(spl)m*β-MS2 foci and condensed Mam hubs in salivary gland nuclei. Aligned Images were averaged and intensities plotted respect to time. For further quantifications, pixel intensities were binned using a 10 x 10 grid. Scale bars represent 5 μm. **(B)** Boxplots with maximum levels and standard deviations of binned Mam intensities within *E(spl)-C* in non-transcribing and transcribing nuclei. **(C)** Heatmap of cross-correlation between paired *E(spl)m*β–MS2 and Mam tracks. Transcription onset is centered (0) and time-lags were applied to Mam tracks as indicated and the correlation calculated, turbo shading indicates strength of correlation.

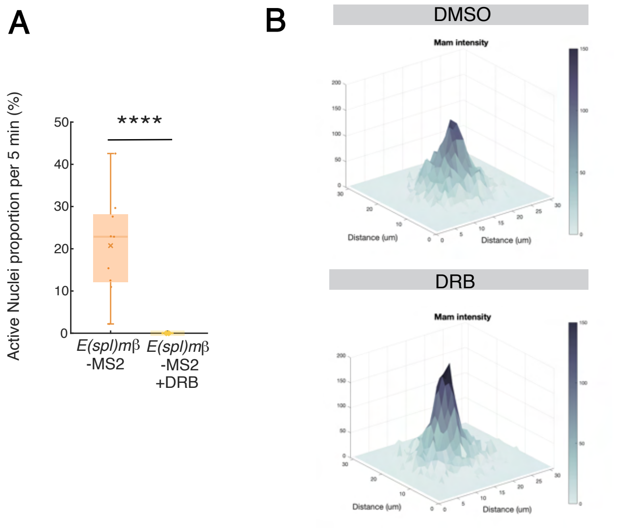


**Figure S5: Transcription inhibition with DRB stabilizes Mam hub.**

**(A-B)** Effect of DRB treatment on transcription and Mam enrichment in follicle cells. **(A)** Average proportions of transcribing nuclei using *E(spl)m*β-MS2 from control (DMSO, n=10) and DRB treated (DRB, n=10) stage 6 egg chambers. **(B)** 3D plot (left panels) of Mam mean intensity at *E(spl)-C* in control (DMSO, n=363) and DRB treated (DRB, n=388) tissues as in A, with intensities partitioned into quartiles (right panels).

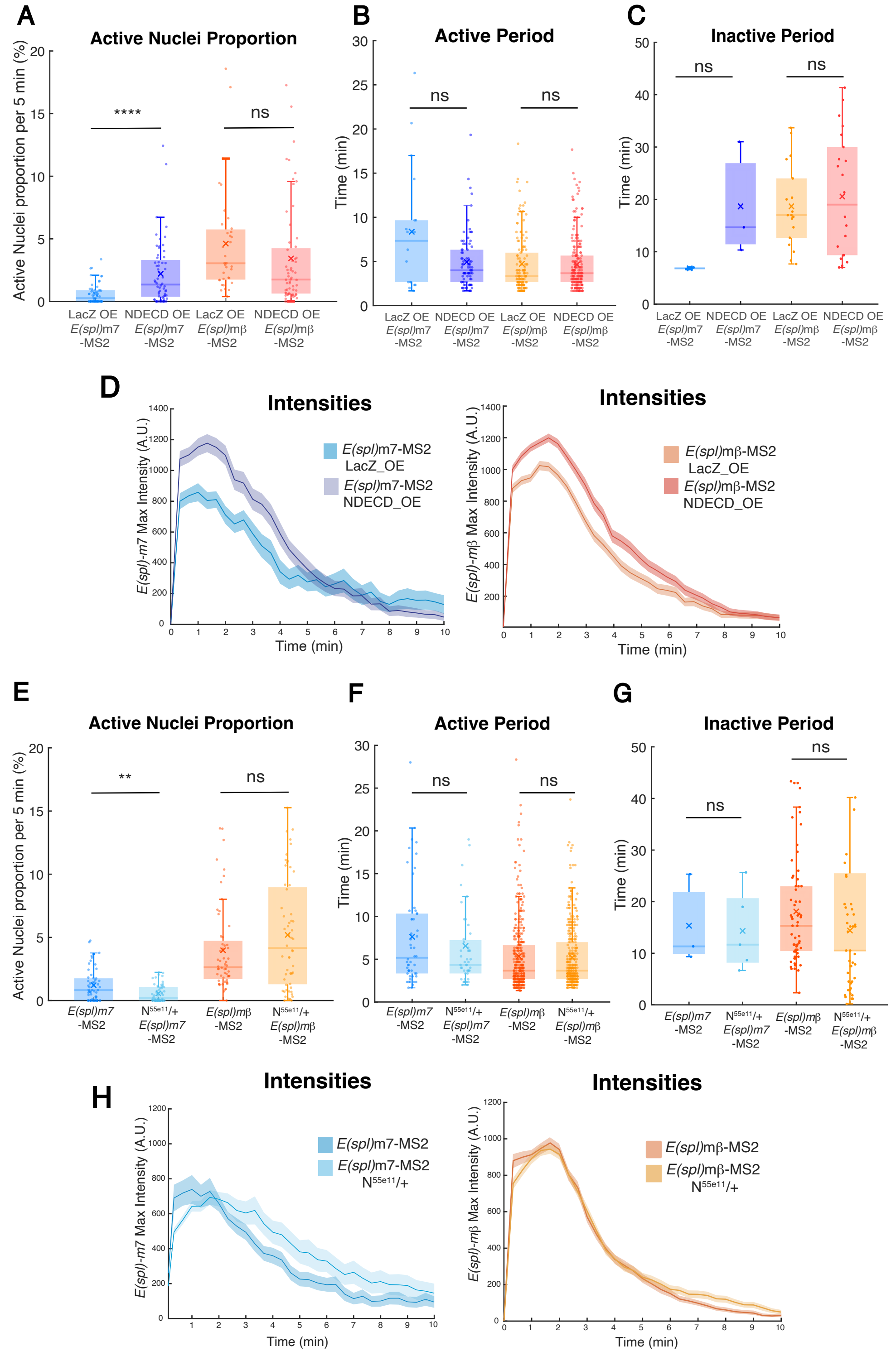


**Fig. S6. smFISH experiments confirm changes of transcription output when Notch dose is altered.**

**(A)** Boxplots of active nuclei proportions from live imaging of *E(spl)m7*-MS2 and *E(spl)m*β-MS2 in control and mild *N*Δ*ECD* overexpression. Duration of active periods are similar for all conditions as indicated. **(B)** Duration of active periods for all conditions.**(C)**Duration of inactive periods for all conditions as indicated. **(D)** Transcription amplitude profiles; mean track intensities from *E(spl)m7*-MS2 (blue, left) and *E(spl)m*β -MS2 (orange, right) in control (mid-blue/orange) and Notch overexpression (dark blue/orange) conditions. **(E)** Boxplots of active nuclei proportions from live imaging of *E(spl)m7*-MS2 and *E(spl)m*β-MS2 in control and *N55e11/+*. **(F)** Duration of active periods for all conditions. **(G)** Duration of inactive periods for all conditions as indicated. **(H)** Transcription amplitude profiles; mean track intensities from *E(spl)m7*-MS2 (blue, left) and *E(spl)m*β-MS2 (orange, right) in control (mid-blue/orange) and Notch heterozygotes (light blue/orange) conditions. Boxplots indicate median, with 25–75 quartiles; error bars are SD. In D,I graphs, SEM is represented by shading.

**A**

| | *E(spl)m7* | *E(spl)mβ* |
|---|---|---|
| **RNA Pol II speed (bp/s)** | 45 | 45 |
| **Time Resolution (s)** | 20 | 20 |
| **Min Pol II distance (bp)** | 30 | 30 |
| **mRNA length pre MS2 (bp)** | 679 | 0 |
| **MS2 length (bp)** | 1330 | 1330 |
| **mRNA length post MS2 (bp)** | 3128 | 4106 |

**B**

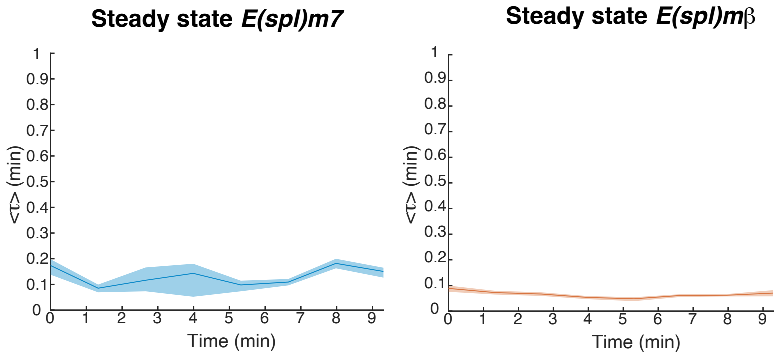


**Figure S7: Parameters and Steady state confirmation for modelling.**

**(A)** Parameters feeded to the BURSTDECONV model for *E(spl)m7*-MS2 and*E(spl)m*β-MS2. **(B)** Average interval between successive Pol II initiation events (<τ>) through a 4 frames time window for *E(spl)m7*-MS2 and*E(spl)m*β-MS2 confirmed that steady state has been reached stage 6.

**Supplementary Table S1: Details of Drosophila lines used.**

| Line / Element | Genotype (FlyBase Nomenclature) | Source |
|---|---|---|
| *Delta::mScarlet-I* | *TI{TI}Delta[mScarlet-I]* | [72] |
| Locus tag *(E(spl)-C locus + ParB1)* | *TI{P{w[+]=intA}3xP3-RFP.attP}E(spl)mδ-HLH, P{w[+]=UAS-ParA::mCherry}attP86Fb / TM6B* | [16] |
| *GFP::CSL* | *M{ w[+]=Su(H)::EGFP}attP86Fb* | [15] |
| *Halo::CSL* | *M{ w[+]=Su(H)::Halo}attP86F* | [15] |
| *GFP::Hairless* | *M{ w[+]=H.WT.EGFP}51D* | [16] |
| *Halo::Mam* | *TI{TI}mamRA[Halo]* | [15] |
| *GFP::Mam* | *TI{TI}mamRA[sfGFP]* | [15] |
| *Sox14::GFP* | *w[1118]; PBac{y[+mDint2] w[+mC]=Sox14-GFP.FPTB}* | BDSC #55842 |
| *E(spl)m7-MS2* | *TI{24xMS2-lacZ-SV40}E(spl)m7-HLH* | This paper |
| *E(spl)mβ-MS2* | *TI{24xMS2-lacZ-SV40}E(spl)mβ-HLH* | [72] |
| *hsp83-MCP::GFP* | *P{w[+mC]=Hsp83-MCP-GFP}3* | BDSC #7280 |
| *His2Av::RFP* | *P{His2Av-mRFP}* | BDSC #23650 |
| *UAS-NΔECD* | *P{w[+]=UAS-NotchΔECD}* | [16] |
| *UAS-NΔECD::mCherry* | *P{w[+]=UAS-NotchΔECD:*mCherry *}* | This paper |
| *tj-GAL4* | *w[*], P{w[+]=GawB}Tj* | St Johnston lab |
| *tj-GAL4::tub-Gal80*$^{ts}$ | *w[*], P{w[+]=GawB}Tj , P{w[+]=tubP-GAL80[ts]}* | St Johnston lab |
| *UAS-LacZ* | *P{w[+]=UAS-nls-LacZ}* | Brand lab |
| *hs-Flp* | *P{ry[+t7.2]=hsFLP}1, y[1] w[1118]; Dr[1]/TM3, Sb[1]* | BDSC #26902 |
| *Tub>FRT.STOP.FRT>GAL4::UAS-mTandemTomato* | *w[1118]* ; P{*tubP[FRT.CD2-Stop] GawB}*, 10x*UAS-IVS-myr::tdTom in attP40* / CyO,P{w[+mC]=Dfd-GMRYFP}2 | Van den Ameele lab |
| $N^{55e11}$ *FRT19A* | *w[1118] N[55e11] P{ry[+t7.2]=neoFRT}19A* | BDSC #28813 |
| *yw* | *Y[1] w[1118]* | BDSC #6598 |

**Supplementary Table S2: Detailed genotypes for each Figure.**

| Figure | X chromosome | II chromosome | III chromosome |
|---|---|---|---|
| Figure 1B | *w* | *Sco/CyO* | *Delta-*$^{mScarlet\text{-}I}$ |
| Figure 1D | *w* | *tj-Gal4* | *Halo::CSL, E(spl)-C[m∂ intA}, UAS-ParA::mCherry* |
| Figure 1D | *w* | *tj-Gal4/GFP::Hairless* | *E(spl)-C[m∂ intA}, UAS-ParA::mCherry* |
| Figure 1D | *w* | *tj-Gal4/Halo::Mam* | *E(spl)-C[m∂ intA}, UAS-ParA::mCherry* |
| Figure 1E | As figure 1D | | |
| Figure 2A,B | *yw* | | |
| Figure 2D | *w* | *hsp83-MCP::GFP* | *E(spl)mβ-MS2/ His2Av-RFP* |
| Figure 2E,F,G,H | *w* | *hsp83-MCP::GFP* | *E(spl)mβ-MS2/ His2Av-RFP* |
| Figure 2E,F,G,H | *w* | *hsp83-MCP::GFP* | *E(spl)m7-MS2/ His2Av-RFP* |
| Figure 3A | *w* | *tj-Gal4/Halo::Mam* | *E(spl)-C[m∂ intA}, UAS-ParA::mCherry* |
| Figure 3B,C,F | *w* | *hsp83-MCP::GFP, tj-Gal4/Halo::Mam* | *E(spl)m7-MS2/ E(spl)-C[m∂ intA}, UAS-ParA::mCherry* |
| Figure 3D,E,G | *w* | *hsp83-MCP::GFP, tj-Gal4/Halo::Mam* | *E(spl)mβ -MS2/ E(spl)-C[m∂ intA}, UAS-ParA::mCherry* |
| Figure 4 | *1151-Gal4* | *hsp83-MCP::GFP /Halo::Mam* | *E(spl)mβ -MS2/ UAS-NΔECD* |
| Figure 5A | *w* | *tj-Gal4/Halo::Mam* | *E(spl)-C[m∂ intA}, UAS-ParA::mCherry* |
| Figure 5B,C,E | *1151-Gal4* | *hsp83-MCP::GFP /Halo::Mam* | *E(spl)mβ -MS2/ UAS-NΔECD* |
| Figure 6A,B | *w* | *tj-GAL4::tub-Gal80*$^{ts}$*/Halo::Mam* | *E(spl)-C[m∂ intA}, UAS-ParB::GFP/ UAS-NΔECD::mCherry* |
| Figure 6A,B | *w* | *tj-GAL4::tub-Gal80*$^{ts}$*/Halo::Mam* | *E(spl)-C[m∂ intA}, UAS-ParB::GFP/ UAS-LacZ* |
| Figure 6C,D | *w* | *tj-GAL4::tub-Gal80*$^{ts}$ | *UAS-NΔECD::mCherry* |
| Figure 6C,D | *w* | *tj-GAL4::tub-Gal80*$^{ts}$ | *UAS-LacZ* |
| Figure 6F,G | *N*$^{55e11}$ *FRT19A* | *tj-Gal4/Halo::Mam* | *E(spl)-C[m∂ intA}, UAS-ParA::mCherry* |

| Figure | X chromosome | II chromosome | III chromosome |
|---|---|---|---|
| Figure 6F,G | *w* | *tj-Gal4/Halo::Mam* | *E(spl)-C[m∂ intA}, UAS-ParA::mCherry* |
| Figure 6H,I | *$N^{55e11}$ FRT19A* | | |
| Figure 6H,I | yw | | |
| Figure S1A | *w* | *tj-Gal4* | *Halo::CSL, E(spl)-C[m∂ intA}, UAS-ParA::mCherry* |
| Figure S1B | *w* | *tj-Gal4/Halo::Mam* | *E(spl)-C[m∂ intA}, UAS-ParA::mCherry* |
| Figure S1C | *w* | *tj-Gal4/Sox14::GFP* | *E(spl)-C[m∂ intA}, UAS-ParA::mCherry* |
| Figure S1D | *w* | | *GFP::CSL* |
| Figure S1D | *w* | *GFP::Hairless* | |
| Figure S1D | *w* | *GFP::Mam* | |
| Figure S3A | *w* | *hsp83-MCP::GFP, tj-Gal4/Halo::Mam* | *E(spl)mβ -MS2/ E(spl)-C[m∂ intA}, UAS-ParA::mCherry* |
| Figure S3B | *w* | *tj-Gal4/Halo::Mam* | *E(spl)-C[m∂ intA}, UAS-ParA::mCherry* |
| Figure S4 | *1151-Gal4* | *hsp83-MCP::GFP /Halo::Mam* | *E(spl)mβ -MS2/ UAS-NΔECD* |
| Figure S5A, B,C,D | *w* | *tj-Gal4/Halo::Mam* | *E(spl)-C[m∂ intA}, UAS-ParA::mCherry* |
| Figure S5E,F | *1151-Gal4* | *hsp83-MCP::GFP /Halo::Mam* | *E(spl)mβ -MS2/ UAS-NΔECD* |
| Figure S6A,B,C,D | *hs-Flp* | *Tub>FRT.STOP.FRT>GAL4::UAS-mTandemTomato/ hsp83-MCP::GFP* | *E(spl)m7 -MS2/ UAS-NΔECD::mCherry* |
| Figure S6A,B,C,D | *hs-Flp* | *Tub>FRT.STOP.FRT>GAL4::UAS-mTandemTomato/ hsp83-MCP::GFP* | *E(spl)m7 -MS2/ UAS-LacZ* |
| Figure S6A,B,C,D | *hs-Flp* | *Tub>FRT.STOP.FRT>GAL4::UAS-mTandemTomato/ hsp83-MCP::GFP* | *E(spl)mβ -MS2/ UAS-NΔECD::mCherry* |
| Figure S6A,B,C,D | *hs-Flp* | *Tub>FRT.STOP.FRT>GAL4::UAS-mTandemTomato/ hsp83-MCP::GFP* | *E(spl)mβ -MS2/ UAS-LacZ* |
| Figure S6E,F,G,H | *$N^{55e11}$ FRT19A* | *hsp83-MCP::GFP* | *E(spl)m7-MS2/ His2Av-RFP* |

| Figure | X chromosome | II chromosome | III chromosome |
|---|---|---|---|
| Figure S6E,F,G,H | w | *hsp83-MCP::GFP* | *E(spl)m7-MS2/ His2Av-RFP* |
| Figure S6E,F,G,H | $N^{55e11}$ *FRT19A* | *hsp83-MCP::GFP* | *E(spl)mβ-MS2/ His2Av-RFP* |
| Figure S6E,F,G,H | w | *hsp83-MCP::GFP* | *E(spl)mβ-MS2/ His2Av-RFP* |

**Supplementary Table S3: results of statistical analysis of data presented in Figures**

| Figure | Experiment | n | p Value |
|---|---|---|---|
| Figure 1D | CSL. St 5, 6, 7, 8, 9, 10 | n=221, 137 377,137,259, 158 | |
| Figure 1D | Mam. St 5, 6, 7, 8, 9, 10 | n=92, 141, 173, 171, 287, 219 | |
| Figure 1D | Hairless. St 5, 6, 7, 8, 9, 10 | n= 94, 186, 241, 158, 248, 207 | |
| Figure S1A | CSL. St 5, 6, 7, 8, 9, 10 | n= 36, 61, 61, 61, 61, 61 | P=0 |
| Figure S1B | Hairless. St 5, 6, 7, 8, 9, 10 | n=61, 59, 61, 61, 61, 61 | P= 4.664e-11 |
| Figure S1C | Mam. St 5, 6, 7, 8, 9, 10 | n=46, 53, 37, 37, 57,57 | P=0 |
| Figure 2B | Active nuclei *E(spl)m7 vs E(spl)mβ* stage 6 | n=20, 18 | p=4,5e-10 |
| Figure 2G | Active nuclei *E(spl)m7-MS2 vs E(spl)mβ–MS2* | n=57, 70 | p=3.16e-9 |
| Figure 2H | Active periods *E(spl)m7-MS2 vs E(spl)mβ–MS2* | n=54,342 | p = 0.00114 |
| Figure 2I | Inactive periods *E(spl)m7-MS2 vs E(spl)mβ–MS2* | n=3,63 | P=0.7464 |
| Figure S2A | Active nuclei *E(spl)m7 vs E(spl)mβ* stages 5,6 & 7 | n=20, 16, 20,18,24,21 | |
| Figure S2D | Inactive periods *E(spl)m7-MS2 vs E(spl)mβ–MS2* | n=3,63 | P=0.7464 |
| Figure 3A | 3D Mam stage 6 | n=141. 3 e.c. | |
| Figure 3C | Mam with *E(spl)m7-MS2* in transcribing cells | n=12. 7 e.c. | |

| Figure | Experiment | n | p Value |
|---|---|---|---|
| Figure 3D | Mam with *E(spl)mβ*–MS2 in transcribing cells | n=15. 5 e.c. | |
| Figure 3E | Mam with *E(spl)mβ*–MS2 in no transcribing cells | n=7. 4 e.c. | |
| Figure 3F | Cross-correlation Mam with *E(spl)m7-MS2* | n=12 | On figure. |
| Figure 3G | Cross-correlation Mam with *E(spl)mβ*–MS2 | n=15 | On figure. |
| Figure S3B | Quartiles Mam stage 6 | n=141. 3 e.c. | |
| Figure S3C | Correlation Mam with *E(spl)m7-MS2* slope | n=12 | On figure. |
| Figure S3C | Correlation Mam with *E(spl)mβ–MS2* slope | n=15 | On figure. |
| Figure 4C | Mam with *E(spl)mβ*–MS2 in transcribing cells. SGs | n=20 | |
| Figure 4D | Cross-correlation Average Mam with *E(spl)mβ*–MS2 | n=20, (435 regions) | On figure. |
| Figure 4E | 3D plot transcribing and no transcribing cells. SGs. | n=7, 10 | |
| Figure 4F | Mam autocorrelation in transcribing and no transcribing cells. SGs. | n=7,10 | On figure. |
| Figure S4B | Levels of Mam and SD in no transcribing and transcribing cells. SGs. | n=10 (276 regions), 7 (435 regions) | |
| Figure S4C | Cross-correlation Average Mam with *E(spl)mβ*–MS2. | n=20, (435 regions) | On figure. |
| Figure 5A | 3D Mam in DMSO vs.Triptolide | n= 531, 540. 7, 6 e.c. | |
| Figure 5B | Active nuclei proportion of *E(spl)m7-MS2* before vs after triptolide | n=3, 3 | p= 4.22e-5 |
| Figure 5C | 3D Mam and *E(spl)mβ*–MS2 | n=9 (327 regions), 6 (345 regions) | |

| Figure | Experiment | n | p Value |
|---|---|---|---|
| | transcription in DMSO vs.Triptolide | | |
| Figure 5D | 3D Mam and *E(spl)mβ*–MS2 in DMSO vs. A485 | n=9 (327 regions), 6 (191 regions) | |
| Figure S5A | Active nuclei proportion of *E(spl)mβ–MS2* in control vs DRB. | n=10, 10 | p= 8.86e-5 |
| Figure S5B | 3D Mam in DMSO vs.DRB | n=363, 388 | |
| Figure 6B | 3D Mam in Control and *NΔECD* | n=240, 279 from 5, 6 e.c. | |
| Figure 6D | Active Nuclei proportion *E(spl)m7* control *vs NΔECD* | n= 4, 5 | p= 0.234 |
| Figure 6F | Active Nuclei proportion *E(spl)mβ* control *vs NΔECD* | n= 4, 5 | p=0.1905 |
| Figure 6F | 3D Mam in Control and $N^{55e11}$/+ | n=137, 76 | |
| Figure 6H | Active Nuclei proportion *E(spl)m7* control *vs* $N^{55e11}$/+ | n=15, 9 | p=0.455 |
| Figure 6H | Active Nuclei proportion *E(spl)mβ* control *vs* $N^{55e11}$/+ | n=11,9 | p=0.062 |
| Figure S6A | Active Nuclei proportion *E(spl)m7-MS2* control *vs NΔECD* | n= 5 , 5 e.c. | p=7.88e-6 |
| Figure S6A | Active Nuclei proportion *E(spl)mβ–MS2* control *vs NΔECD* | n=3, 5 e.c. | p=0.18 |
| Figure S6B | Active periods *E(spl)m7-MS2* control *vs NΔECD* | n= 18, 84 | p= 0.1185 |
| Figure S6B | Active periods *E(spl)mβ–MS2* control *vs NΔECD* | n= 134, 211 | p = 0.802 |
| Figure S6C | Inactive periods *E(spl)m7-MS2* control *vs NΔECD* | n=2, 3 | p = 0.2 |

| Figure | Experiment | n | p Value |
|---|---|---|---|
| Figure S6C | Inactive periods *E(spl)mβ–MS2* control *vs NΔECD* | n=18, 22 | p = 0.92 |
| Figure S6D | Intensities *E(spl)m7-MS2* control *vs NΔECD* | n=64 tracks from 5 e.c., 85 tracks from 5 e.c. | |
| Figure S6D | Intensities *E(spl)mβ–MS2* control *vs NΔECD* | n=155 tracks from 3 e.c., 231 tracks from 5 e.c. | |
| Figure S6E | Active Nuclei proportion *E(spl)m7-*MS2 control *vs* $N^{55e11}$/+ | n=5, 6 e.c. | p = 7.88e-6 |
| Figure S6E | Active Nuclei proportion *E(spl)mβ–MS2* control *vs* $N^{55e11}$/+ | n= 6, 5 e.c | p = 0.184 |
| Figure S6F | Active periods *E(spl)m7-MS2* control *vs* $N^{55e11}$/+ | n=52, 47 | p = 0.381 |
| Figure S6F | Active periods *E(spl)mβ–MS2* control *vs* $N^{55e11}$/+ | n=340, 326 | p = 0.649 |
| Figure S6G | Inactive periods *E(spl)m7-MS2* control *vs* $N^{55e11}$/+ | n=3, 5 | p = 4.215e-8 |
| Figure S6G | Inactive periods *E(spl)mβ–MS2* control *vs* $N^{55e11}$/+ | n=63, 45 | p = 0.227 |
| Figure S6H | Intensities *E(spl)m7-MS2* control *vs* $N^{55e11}$/+ | n= 66 tracks from 5 e.c., 35 tracks from 6e.c. | |
| Figure S6H | Intensities *E(spl)mβ–MS2* control *vs* $N^{55e11}$/+ | n=443 from 6 e.c., 318 from 5 e.c. | |
